# Sugar Signaling Induces Dynamic Changes During Meristem Development to Regulate Flowering in Arabidopsis

**DOI:** 10.1101/2021.04.12.439483

**Authors:** Magdalena Musialak-Lange, Katharina Fiddeke, Annika Franke, Friedrich Kragler, Christin Abel, Vanessa Wahl

## Abstract

Aerial parts of plants originate from pluripotent stem cells in the shoot apical meristem. Their population is maintained via the maintenance regulators *WUSCHEL* and *CLAVATA3* in a negative feed-back loop. Meristem size is dynamic and undergoes a more than 2-fold expansion upon the transition to reproductive growth. The mechanism controlling this doming is largely unknown, but coinciding increased trehalose 6-phosphate and changed meristem size in overexpressing or knockdown lines of *TREHALOSE PHOSPHATE SYNTHASE1* suggest a participation of sugar signaling. Here we show that *TREHALOSE PHOSPHATE PHOSPHATASEJ* is directly regulated by WUSCHEL. Plants with reduced levels of *TREHALOSE PHOSPHATE PHOSPHATASEJ* in the outer meristem layer are flowering early and its reduction in the late flowering *clavata3* mutant, restores wild-type flowering. This is caused by a reduction of mature miR156 and increased expression of *SQUAMOSA PROMOTER-BINDING PROTEIN-LIKE* genes. We demonstrate that these are important for age pathway-induced flowering, in a negative feed-back loop with WUSCHEL downstream of the trehalose 6-phosphate pathway. In summary, our findings demonstrate a dynamic feed-back regulation between central maintenance and flowering time regulators with sugar signaling.

**Synopsis:**
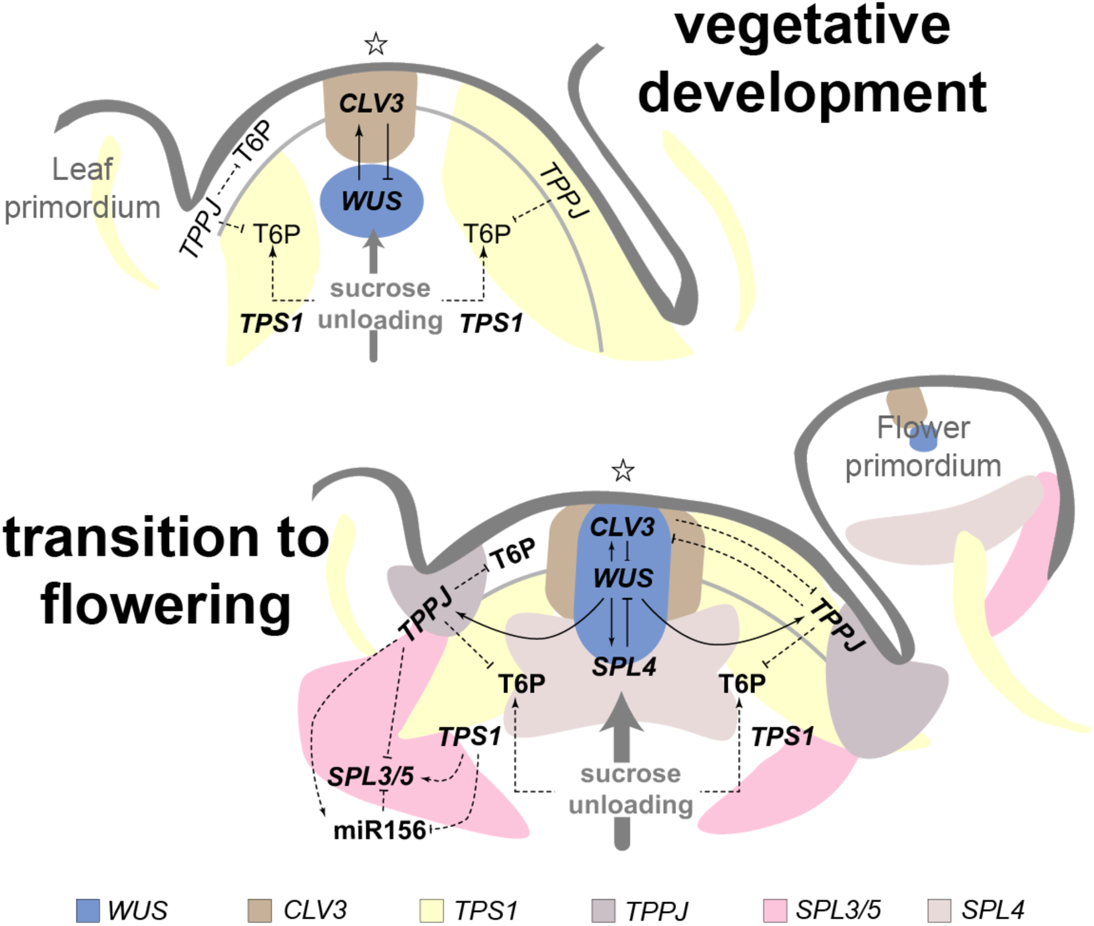
Schematic illustration of dynamic feedback-regulations between sugar signaling, meristem maintenance and the age pathway of the flowering network in the transition SAM as compared to the vegetative SAM. During vegetative growth SAM maintenance is controlled by spatially separated WUS and CLV3 expression in a negative feedback-loop. At floral transition increased activity of the T6P pathway in the SAM uncouples this regulation. This results in an unreported special relocation of WUS expression and involves a transient negative feedback-loop between WUS and SPL4.

## Introduction

All above ground plant organs such as leaves and flowers originate from the shoot apical meristem (SAM). The *Arabidopsis thaliana* (Arabidopsis) SAM is organized in three clonally distinct cell layers: the outer mono cell layers, L1 and L2, and the several cell files comprising L3. Functionally, the SAM consist of a central zone (CZ), containing undifferentiated cells with a self-renewing potential, an organizing centre (OC), inducing and maintaining an adequate number of cells in the CZ, a peripheral zone (PZ) which receives cells from the CZ that are differentiating and incorporated into leaf or flower primordia, and a rib zone (RZ) producing the stem (Pfeiffer *et al*, 2017). Communicating position-dependent properties among cells within and between the different zones is crucial for SAM function. Two non-cell autonomously acting factors regulate each other in a negative feed-back loop. The homeodomain transcription factor *WUSCHEL* (*WUS*) is expressed in the OC, and the signalling peptide *CLAVATA3* (*CLV3*) in the CZ. Thus, mobile WUS transcription factor, secreted from the OC, activates expression of *CLV3* in the CZ above, while CLV3 peptide restricts the *WUS* expression domain to the OC. Hence, *WUS* and *CLV3* expression domains are considered mutually exclusive and their negative feed-back regulation as very robust, which can only be modulated within tight limits (Pfeiffer *et al*., 2017).

Development as a whole and SAM maintenance in particular demands continuous cross talk between its regulatory processes and the available resources. SAM maintenance and flowering time are important determinants of crop yield. During floral transition cell division is activated and the SAM undergoes a dramatic increase in size known as doming (Jacqmard *et al*, 2003; Kwiatkowska, 2008) (Figure 1A) and marks the time between the vegetative stage (production of leaves) and the reproductive stage (formation of flowers). Since this transition requires a massive reorganization of organ development and sufficient energy, it is tightly controlled by environmental conditions and availability of nutrients (Andres & Coupland, 2012; Olas *et al*, 2019; Ponnu *et al*, 2020; Srikanth & Schmid, 2011; Wahl *et al*, 2013). Only recently the early cellular changes were associated with floral regulators that control GA biosynthesis (Kinoshita *et al*, 2020). However, the knowledge on the mechanisms controlling doming is still scarce.

**Figure 1.**
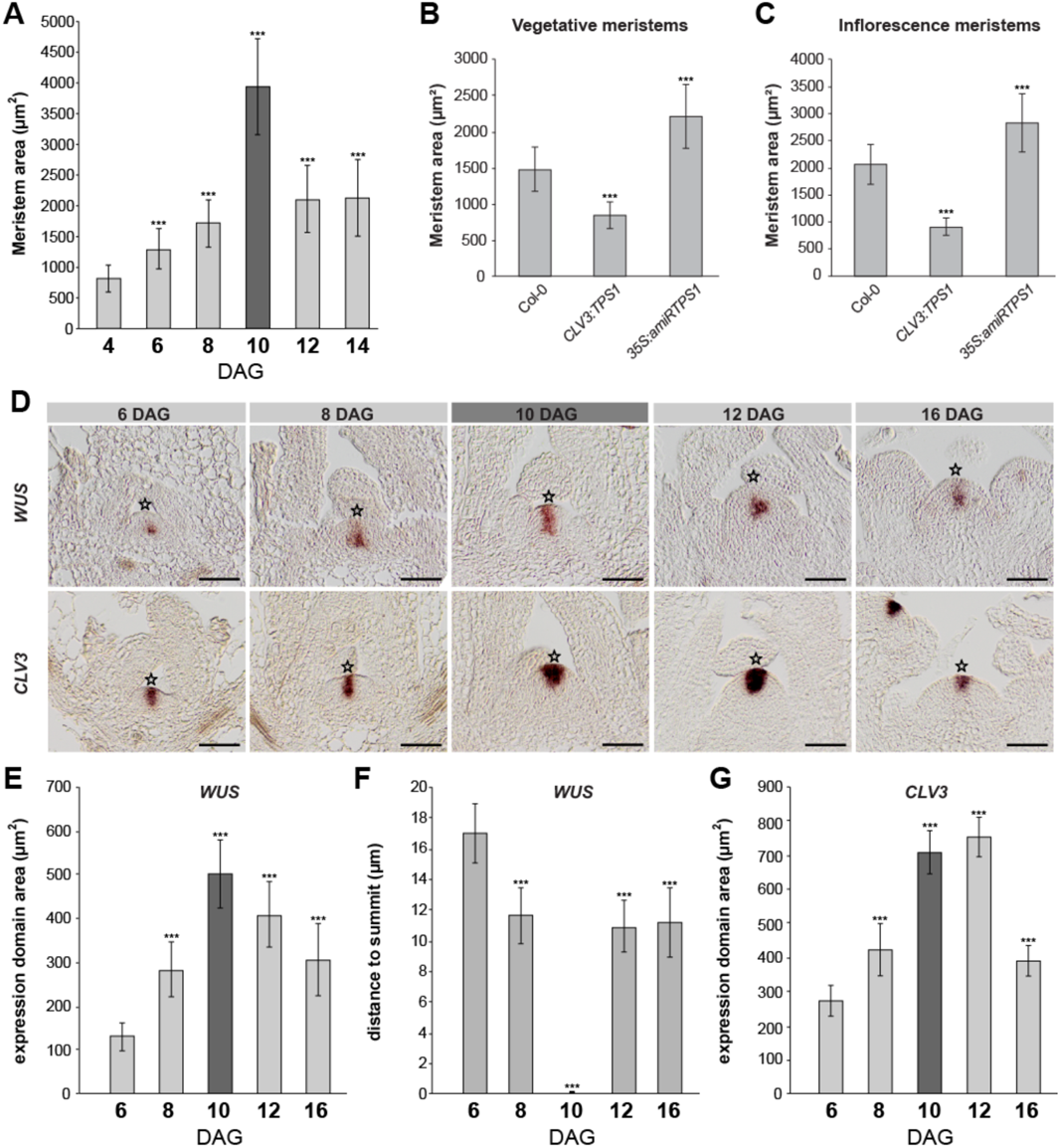
The T6P pathway impacts SAM size during development. (*A*) SAM area throughout development. n=15 (*B*) Vegetative and (*C*) inflorescence SAM size of *CLV3:TPS1* and *35S:amiRTPS1* lines. n=10. (*D*) *WUS* and *CLV3* expression by RNA *in situ* hybridization in vegetative (6 and 8 DAG), transition (10 DAG, marked dark grey) and inflorescence SAMs (12 and 16 DAG) of LD-grown Col-0 plants. (*E*) *WUS* expression domain sizes, (*F*) *WUS* expression domain distance to SAM summit, and (*G*) *CLV3* expression domain area, in vegetative, transition (dark grey) and inflorescence SAMs of LD-grown wild-type plants. n>10. Error bars denote s.d.; significance was calculated based on a Student’s *t*-test, ***P<0.001. Star indicates SAM summit. Scale bars 25µm.

In plants, the sugar phosphate, trehalose 6-phosphate (T6P), it is synthetized from UDP-glucose and glucose 6-phosphate via TREHALOSE PHOSPHATE SYNTHASE (TPS) and is converted to trehalose by TREHALOSE PHOSPHATE PHOSPHATESES (TPP) (Fichtner & Lunn, 2021). T6P serves as a signal for sucrose availability, which is conveyed to downstream metabolic and growth responses through still largely unknown mechanisms (Fichtner & Lunn, 2021). In Arabidopsis sugar signalling affects the vegetative phase change, the transition between the formation of juvenile and adult leaves, via the T6P pathway (Ponnu *et al*., 2020; Yang *et al*, 2013; Yu *et al*, 2013). In addition, the T6P pathway is involved in shoot branching (Fichtner *et al*, 2020), and induces flowering via regulating key flowering genes in leaves and at the SAM (Wahl *et al*., 2013). In leaves, *TPS1*, coding for the T6P synthesizing enzyme, is necessary and sufficient to induce *FLOWERING LOCUS T* (*FT*), the florigen, connecting the photoperiod signal with a physiological signal. In addition, the T6P pathway affects the age-dependent microRNA156 (miR156)/*SQUAMOSA PROMOTER-BINDING PROTEIN-LIKE* (*SPL*) module at the SAM (Wahl *et al*., 2013). Increased T6P levels coincide with doming (Wahl *et al*., 2013), and a spatial change of the *WUS* expression domain. This suggests a participation of sugar signaling in regulating SAM dynamics through transient uncoupling of the negative feed-back regulation between WUS and CLV3. Here we show that WUS directly regulates *TPPJ*. A late flowering phenotype of *clv3* is caused by increased levels of mature miR156 and decreased expression of *SPL* genes and is restored when *TPPJ* is downregulated in the outer meristem layer. We further demonstrate a negative feed-back loop with WUS and SPL4 downstream of the T6P pathway. In summary, we provide evidence for a dynamic feed-back regulation between central meristem maintenance and flowering time regulators with sugar signaling.

## Results and Discussion

### The T6P pathway affects SAM size

To assess whether the morphological changes at the transition SAM (Figure 1A) involve the T6P pathway, we investigated the effect on SAM architecture by decreasing *TPS1* by the means of an artificial microRNA in the SAM proper (*35S:amiRTPS1*; Figure S1) and increasing *TPS1* in the CZ (*CLV3:TPS1*, Figure S2). These plants have smaller and bigger vegetative and reproductive meristems (Figure 1B,C), resulting in smaller and bigger plants, respectively (Wahl *et al*., 2013).

**Figure 2.**
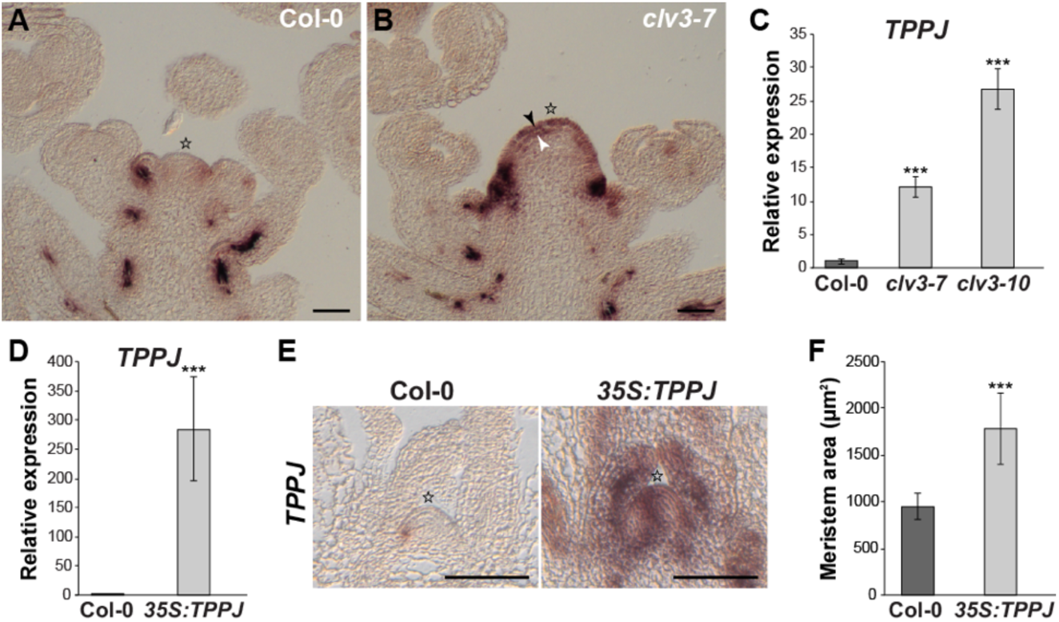
*WUS* and *CLV3* expression in response to changes of *TPPJ* in the SAM. (*A*, *B*) *TPPJ* expression by RNA *in situ* hybridization on longitudinal sections through inflorescence SAMs of (*A*) Col-0, and (*B*) *clv3-7*, and by RT-qPCR in apices collected from (*C*) *clv3-7*, *clv3-10*, and (*D*) *35S:TPPJ* plants. n=3. (*E*) Expression of *TPPJ* by RNA *in situ* hybridization on longitudinal sections through inflorescence SAMs of Col-0 and *35S:TPPJ.* (*F*) SAM size of plants overexpressing *TPPJ*. n=10. Error bars denote s.d.; significance calculated by one-way ANOVA (C,D) and Student’s *t*-test (F), ***P<0.001. Stars indicates SAM summit. Scale bars are 50µm.

*TPS1* knock-out mutants are embryo lethal, likely as a result of insufficient cell cycle activity at torpedo stage of embryogenesis (Gómez *et al*, 2006). Cell cycle genes were also found to be highly induced at transition to flowering (Klepikova *et al*, 2015) confirming earlier findings (Jacqmard *et al*., 2003). In order to test, whether cell proliferation is the cause of the effect observed downstream of the T6P pathway, we used specific probes against transcripts of *HISTONE4, CYCLIN D3;1* and *CYCLIN-DEPENDENT KINASE2;1*, three cell cycle marker genes, for RNA *in situ* hybridization on longitudinal sections through the SAM (Figure S3). The results show more abundant staining indicating the three cell cycle marker genes in the presence of elevated T6P levels and less staining, when T6P levels were reduced. This suggests a higher proliferation rate and an increased number of cells produced to adopt organ-specific cell fates at the periphery as a plausible reason for decreased SAM size of *TPS1* overexpression and *vice versa* increased SAM size of *TPS1* knock-down lines (Figure 1B,C).

**Figure 3.**
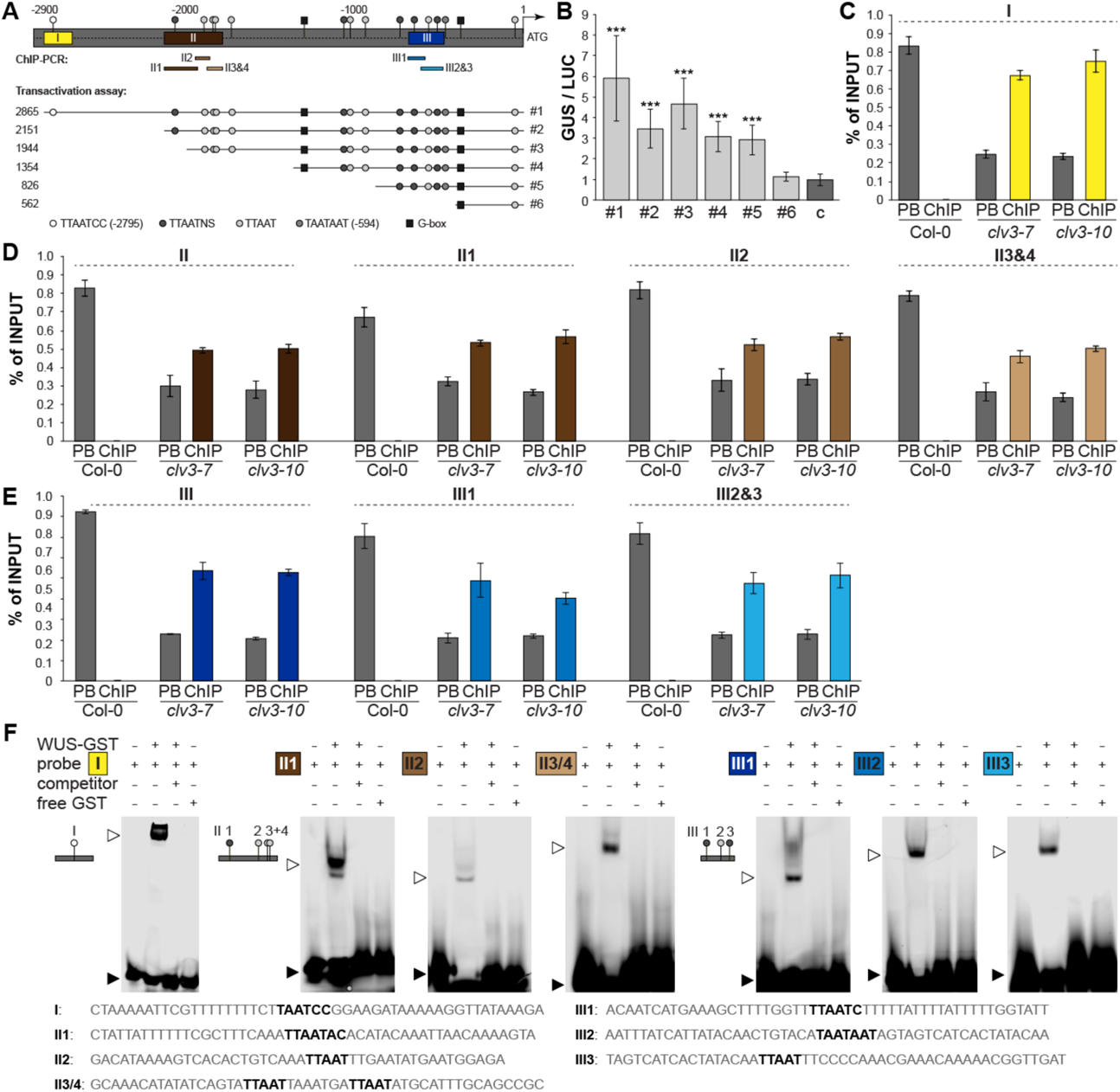
WUS directly regulates TPPJ in the SAM. (*A*) Overview of *TPPJ* 5’ regulatory region with putative *TPPJ^WUS^* sites (grey circles, black boxes), position of ChIP-PCR amplicons corresponding to the results shown in (C-E). Boxes marked with I, II, and III indicate *5’ TPPJ* regions with in total seven confirmed core *TPPJ^WUS^* sites – I: −2795 – −2789 bp, II: −2073 – −1830 bp, and III: −652 – −564 bp. Sequence location and lengths used in (B) indicated with #1-6. (*B*) Protoplast transactivation assay showing activation of the GUS reporter when coupled to the regions indicated in (A), relative to LUC activity. c indicates untransformed control. n=6. (*C*-*E*) Enrichment of (*C*) region I, (*D*) region II, and (*E*) region III as indicated in (A) measured by ChIP-PCR relative to the input. PB – post binding fraction. n=3. (*F*) EMSA for WUS binding to the indicated regions (A, I-III). Shifted band in the presence of WUS protein (open arrow head), non-shifted fraction (closed arrow head). Error bars denote s.d.; significance based on one-way ANOVA, \*\*\**P*<0.001. Scale bars are 50µm.

### Spatial changes of *WUS* expression in a transition SAM

In line with these findings, the floral transition coincides with increased T6P levels (Wahl *et al*., 2013). To assess whether this is related to *WUS* or *CLV3* expression in a wild-type SAM, we analyzed their expression in a time series spanning floral transition (Figure 1D). Throughout the vegetative phase SAM size gradually increases due to rising cell numbers (Figures 1A, S4) (Kinoshita *et al*., 2020), reaching its maximum at floral transition, independent of whether the plants are grown in long days (LD) (Figure 1A) or in short days (SD) followed by a transfer to LD (Figure S5). An enlarging SAM at floral transition correlates with a larger *WUS* expression domain expanding into the CZ and the outer cell layer (L1) (Figure 1D-F).

**Figure 4.**
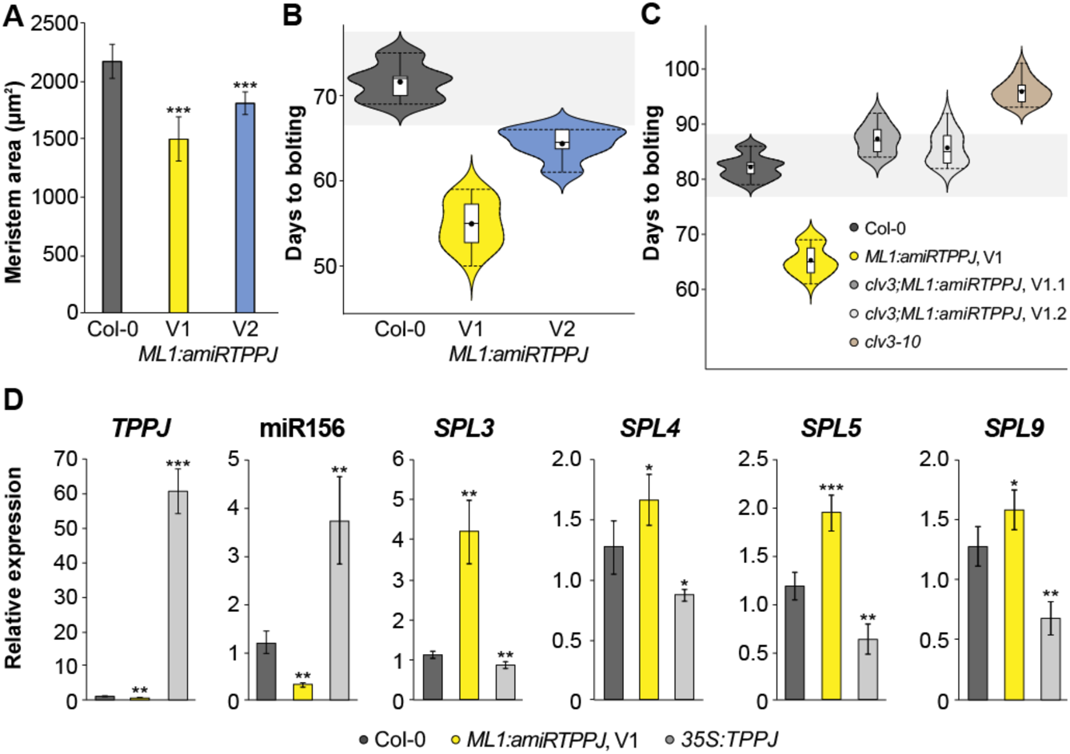
The role of TPPJ in the outer SAM layer. (*A*) Meristem area of Col-0, *ML1:amiRTPPJ* V1 and V2. n=10. (*B*,*C*) Flowering time of (*B*) *ML1:amiRTPPJ* and (*C*) *clv3-10;ML1:amiRTPPJ* shown as days to bolting, relative to the wild type. V1 and V2 indicate two independent versions of artificial microRNAs designed to target *TPPJ* transcript. n=22. (*D*) Relative expression of *SPL* genes in SD-grown *ML1:amiRTPPJ* and *35S:TPPJ* at 40 days after germination. n=4. Error bars denote s.d.; significance calculated based on one-way ANOVA (D) and Student’s *t-*test (A); \**P*<0.05, \*\**P*<0.01, \*\*\**P*<0.001.

**Figure 5.**
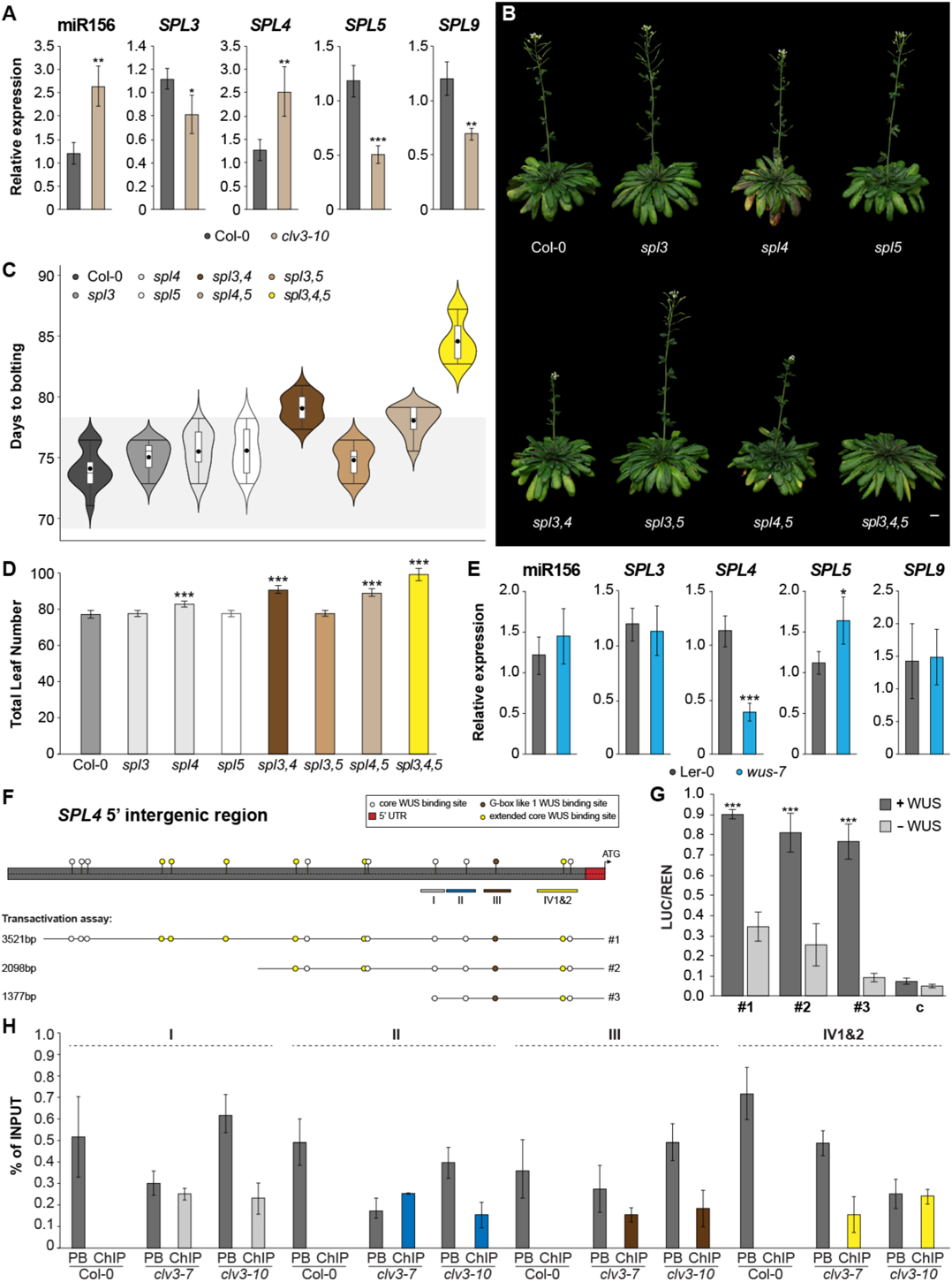
WUS activates *SPL4.* (*A*) Relative expression of mature miRNA and *SPL* genes in SD-grown Col-0 and *clv3-10* apices. n=4 (*B*) Representative pictures of *SPL3-5* single, double and triple CRISPR/*Cas9* deletion mutants in comparison to Col-0 grown in SD. (*C*) Flowering time of deletion mutants displayed in (B) provided as days to bolting and (*D*) total leaf numbers. n=25. (*E*) Relative expression of mature miRNA and *SPL* genes in SD-grown Ler-0 and *wus-7* apices (*F*) Overview of *SPL4* 5’ regulatory region with putative *SPL4^WUS^* sites (grey, yellow and white circles) and position of ChIP-PCR amplicons (grey, blue and yellow boxes) corresponding to the results shown in (G, H). Boxes marked with I, II, III and IV (1&2) indicate *5’ SPL4* regions with in total five core *SPL4^WUS^* sites – I: −1073 – −1068 bp, II: −880 – −875 bp, III: −697 – −691, IV1: −259 – −252 and IV2: −214 – −209 bp. Sequence location and lengths used in (G) indicated with #1-3. (*G*) Protoplast transactivation assay showing activation of the reporter (LUC) when coupled to the regions indicated in (F), relative to REN activity. c indicates vector control. n=3. (*H*) Enrichment of regions I, II, III and IV1&2 as indicated in (F) measured by ChIP-PCR relative to the input in *clv3-7* and *clv3-10* apices. PB – post binding fraction. n=3. Error bars denote s.d.; significance calculated based on one-way ANOVA (A, G) and Student’s *t-*test (D); *P<0.05, \*\**P*<0.01, \*\*\**P*<0.001.

Cytokinin signaling was previously reported to respond to carbon in seedlings (Leibfried *et al*, 2005; Pfeiffer *et al*, 2016; Snipes *et al*, 2018). Its transient occurrence in L1 may thus explain the presence of *WUS* transcript in the outer SAM layers. However, cytokinin levels are not altered in L1 cells as indicated by the synthetic cytokinin reporter *TCSn:GFP* (Zurcher *et al*, 2013) (Figure S6). Further, while *WUS* expands to L1 for a short period (8-10 DAG), the *CLV3* expression domain remains expanded post floral transition (Figure 1D,G). This suggests a transient uncoupling of the negative feed-back regulation between WUS and CLV3, which is re-established at the reproductive SAM (16 DAG), resembling earlier vegetative SAM expression patterns (6 DAG) (Figure 1D).

**Figure 6.**
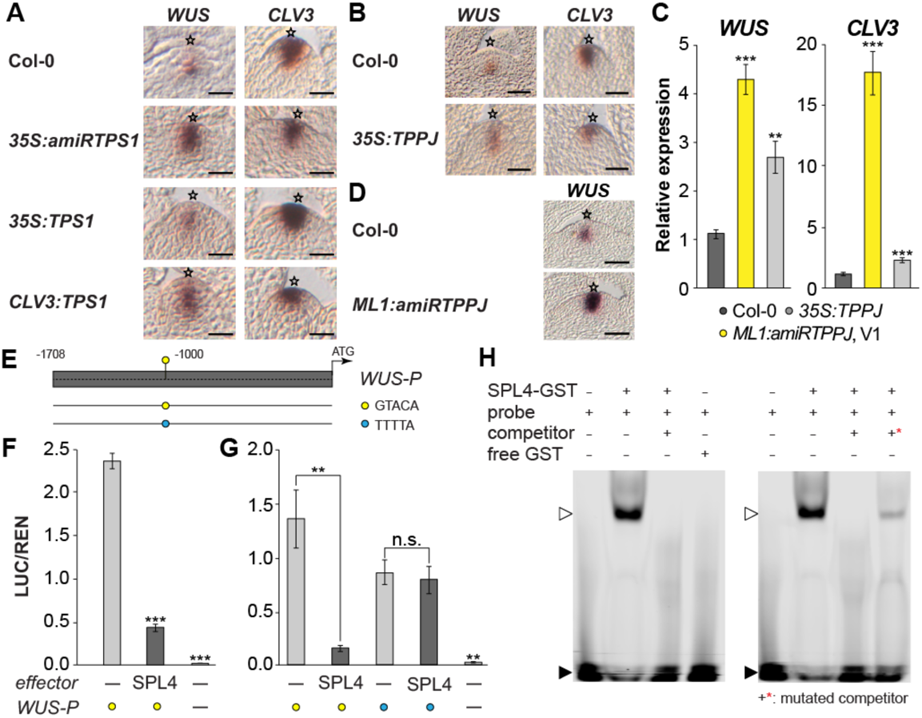
SPL4 negatively feeds back to *WUS.* (*A*,*B*) *WUS* and *CLV3* expression by RNA *in situ* hybridization on (*A*) longitudinal sections through Col-0, *35S:amiRTPS1*, *35S:TPS1* and *CLV3:TPS1* and (*B*) Col-0, and *35S:TPPJ* apices of LD-grown plants. (*C*) Relative expression of *WUS* and *CLV3* in SD-grown Col-0, *35S:TPPJ* and *ML1:amiRTPPJ* V1 apices. n=4 (*D*) RNA *in situ* hybridization on longitudinal sections through Col-0, and *35S:amiRTPPJ* apices. n=3. (*E*) Overview of *WUS* 5’ regulatory region with putative *WUS^SPL^* site corresponding to the results shown in (F,G,H). Grey box indicates *5’ WUS* region with the native and mutated core *WUS^SPL^* site −1067 – −1063 bp (yellow and blue circles, respectively). Sequence location and lengths used in (F,G) indicated as lines below. (*F,G*) Protoplast transactivation assay showing repression of the reporter (LUC) when coupled to the *5’WUS* region indicated in (E), (*F*) relative to REN activity of the native *5’WUS*, and (*G*) the comparison of the native and the mutated *WUS^SPL^* site (yellow and blue circles, respectively), n=3. (*H*) EMSA for SPL4 binding to the *WUS^SPL^* site (E). Shifted band in the presence of SPL4 protein (open arrow head), non-shifted fraction (closed arrow head). Please note the shift in the presence of the mutated competitor (+*), indicating specificity of binding. Error bars denote s.d.; significance calculated based on one-way ANOVA; \*\**P*<0.01, \*\*\**P*<0.001. Scale bars: 25µm. Star indicates SAM summit.

### *TPPJ* is directly regulated by WUS

To understand how the T6P pathway might control this process, we first analyzed expression of the ten genes encoding TREHALOSE PHOSPHATE PHOSPHATASEs (TPP) (Fichtner & Lunn, 2021). Except for *TPPC* and *TPPD*, which are expressed below detection limit of RNA *in situ* hybridization, we found that all are expressed in distinct SAM domains (Figure S7). *TPPJ* strongly increases in the enlarged SAM of *clv3-7* due to an ectopic expression in L1 and L2 (Figures 2A-C; S8, S9). Notably, in *clv3-7* and *clv3-10* mutants high levels of *WUS* in the outer SAM layers (Brand *et al*, 2000) coincide with ectopic expression of *TPPJ* (Figure S9), while *TPPJ* is repressed in a *wus-7* background (Figure S10A). In addition, a previously published data set shows induced levels of *TPPJ* upon DEX induced expression of *WUS* in a *35S:WUS-GR* line (Busch *et al*, 2010) (Figure S10B). To assess whether ectopic expression of *TPPJ* contributes to the enlarged SAM of the *clv3* mutant, we overexpressed *TPPJ* in the wild type (*35S:TPPJ*; Figure 2D,E), which resulted in plants with significantly enlarged SAMs (Figure 2F).

These results imply a direct influence of WUS on *TPPJ*, and is supported by an *in silico* analysis which predicts multiple, canonical WUS binding sites in the sequence upstream of *TPPJ* (*TPPJ*^WUS^; Figure 3A) (Leibfried *et al*., 2005; Lohmann *et al*, 2001; Sloan *et al*, 2020; Yadav *et al*, 2011). To understand if WUS directly controls *TPPJ*, we performed *in vivo* transactivation assays.

These show that WUS activates reporter gene expression when using a 2865bp 5’*TPPJ* sequence, containing 17 putative *TPPJ*^WUS^ sites (Figure 3A). Progressive deletion of this sequence results in a reduction of reporter gene activation, suggesting an additive effect of the individual *TPPJ*^WUS^ sites (#1-6; Figure 3A,B). We confirmed direct binding of WUS to three distinct regions of the *TPPJ* promoter (I, II, and III; Figures 3C-E; S11) by chromatin immunoprecipitation (ChIP) coupled to PCR using a specific antibody against WUS (Figures S11, S12), while all other regions did not indicate binding (Figures S11, S13). We observed enrichment of WUS binding to *TPPJ*^WUS^ sites up to 0.67% of the input DNA in *clv3-7* and 0.75% of the input DNA in *clv3-10* apices, both of which express *WUS* at high levels in comparison to input DNA from wild-type apices, where *WUS* is expressed in only a few cells and input DNA from leaves, with no WUS (Figures 3C-E; S10). Electrophoretic mobility shift assays (EMSA) confirm specific *in vitro* binding of WUS to the investigated sequences (I, II 1-3, III 1-3; Figure 3F; S14). In summary, we demonstrate that seven out of 17 putative *TPPJ*^WUS^ sites are directly targeted by WUS *in vivo*.

### TPPJ controls flowering from the outer SAM layer

A *tppj* mutant with a T-DNA insertion in the 6^th^ exon, which specifically knocks-out *TPPJ* expression (Figure S15), flowers significantly earlier than the wild type (Table S1). To further assess the role of *TPPJ* at the SAM we used an artificial miRNA (amiR) approach to downregulate *TPPJ* (*35S:amiRTPPJ*; Figure S16)(Schwab *et al*, 2006). Plants overexpressing either of two versions of an *amiRTPPJ* (V1, V2) flower significantly earlier in LD and SD (Figure S17; Table S1). *ML1* expression specifically and stably localizes to L1 in all investigated stages (Figure S18). *ML1:amiRTPPJ* reduces *TPPJ* expression in L1 (Figure S19), has a smaller SAM and flowers significantly earlier in LD and SD (Figures 4A,B; S20, S21; Table S1), while the level of *TPPJ* reduction is proportional to the acceleration of flowering (Figures 4B,C; S16; Table S1).

To date there are no reports that mutants in meristem maintenance genes have a flowering phenotype. We found that *clv3* plants are late flowering (Figure 4C; Table S1). However, when *ML1:amiRTPPJ* is introgressed into the *clv3-7* or *clv3-10* background, the late flowering phenotype is restored to wild-type (Figures 4C; S21; Table S1), suggesting that *TPPJ* expression in the outer meristem layers is causal for the late flowering phenotype of *clv3*. Hence, the early flowering of *ML1:amiRTPPJ* in a wild-type background is due to a reduction of *TPPJ* expression in the outer meristem layer in its endogenous expression domain. In addition, other prominent morphological defects of *clv3-10* such as the fasciated stem and enormous inflorescence SAM are visibly reduced in the presence of *ML1:amiRTPPJ* (Figure S21).

### WUS regulates the age pathway through *SPL4*

We previously reported that the T6P pathway influences the age pathway at the SAM (Wahl *et al*., 2013). We therefore next analyzed mature miR156, as well as the expression of *SPL3*, *SPL4*, *SPL5*, *SPL9* and *SPL15*, all associated with floral transition in one or the other way (Hyun *et al*, 2017; Xie *et al*, 2020). We found a strong reduction of mature miR156 levels correlating with decreased *TPPJ* in *ML1:amiRTPPJ*, and corresponding increased expression of *SPL3, SPL4, SPL5*, and *SPL9* (Figure 4D). In line, miR156 was more abundant in *35S:TPPJ*, while the corresponding *SPL*s were downregulated (Figure 4D). Expression of *SPL15*, previously described as an important integrator of plant age into the regulation of floral onset in SD (Hyun *et al*, 2016; Wang *et al*, 2009; Yamaguchi *et al*, 2014), was not differentially expressed in the transgenic lines (Figure S22A).

Uncoupling of the WUS/CLV3 feed-back loop occurs downstream of the T6P pathway and involves age pathway components. Interestingly, a study by Fouracre and Poethig indicates an active pool of miR156 in the SAM, which is essential for early shoot maturation and affects leaf identity. In addition, apices of a *wus-5* mutant had decreased miR156 levels and increased *SPL9* expression, although expression of none of the other SPLs was tested. However, plants with increased miR156 (*35S:miR156a*) have a larger SAM, while those with reduced miR156 (quadruple *mir156a,c,mir157a,c*) have a smaller SAM, and a sextuple *spl2,9,10,11,13,15* increased SAM size. This is accompanied with an increased *WUS* expression in the sextuple mutant, suggesting the potential repression of *WUS* by any or all of the *SPL*s mutated in hextuple plants (Fouracre & Poethig, 2019). Notably, we found miR156 levels significantly increased in *clv3-10* apices (Figure 5A). In response, *SPL3*, *SPL5*, and *SPL9* were reduced supporting the late flowering phenotype of the mutant, while *SPL15* was not affected (Figure S22B). In contrast to what would be expected, we found more *SPL4* transcript in apices of the late flowering *clv3-10* (Figure 5A).

The contribution of SPL9 and SPL15 to the regulation of the floral onset is well established (Hyun *et al*., 2016; Wang *et al*., 2009; Yamaguchi *et al*., 2014), but there is some dispute about whether or not SPL3, SPL4, and SPL5 are important. While overexpression of *SPL3, SPL4*, and *SPL5* induces flowering (Jung *et al*, 2016; Wu & Poethig, 2006), a triple T-DNA insertion mutant was previously reported to not display any flowering phenotype (Xu *et al*, 2016). However, a recent study analyzed a triple deletion mutant generated by CRISPR/*Cas9* in relation to light signaling and also reported on a delay in flowering time (Xie *et al*., 2020). Similarly, we generated individual deletion mutants in the respective SPLs using CRISPR/*Cas9* (Figure S23). As compared to single *spl* mutant and wild-type plants, higher order *spl3*, *spl4* and *spl5* deletion mutants are significantly later flowering in all lines with *spl4* (Figure 5B-D; Table S1). This argues for an important role of *SPL4* in inducing flowering. *SPL4*, similar to *SPL3* and *SPL5,* is induced at the wild-type SAM at floral transition (Schmid *et al*, 2003). It is expressed in the center of the SAM, in a domain overlapping with the WUS protein, while *SPL3*, *SPL5*, *SPL9*, and *SPL15* are expressed at the periphery of the SAM and in the vasculature of young leaves (Daum *et al*, 2014; Hyun *et al*., 2016; Olas *et al*., 2019; Wang *et al*., 2009; Yadav *et al*., 2011). This denotes a direct regulation of *SPL4* by WUS, which would explain increased *SPL4* levels in the *clv3* mutant (Figure 5A) and the previously identified partial miR156-dependent regulation of the *SPL*s by the T6P pathway (Wahl *et al*., 2013). Indeed, we observed reduced expression of *SPL4* in *wus-7* apices, a weak allele with a mutation in the WUS DNA binding domain resulting in smaller but maintained SAM tissue (Graf *et al*, 2010; Lin *et al*, 2016). In contrast, none of the other *SPL*s were reduced (Figures 5E; S22C) and miR156 levels did not change (Figure 5E). This is unexpected as a recent publication demonstrated decreased miR156 levels and increased *SPL9* expression in apices of a *wus-5* mutant (Fouracre & Poethig, 2019). *wus-5* is however a strong *wus* null allele with very little residual SAM tissue (McElver *et al*, 2001; Sonoda *et al*, 2007), which might explain the difference. We identified a larger number of potential *SPL4*^WUS^ sites when compared to the other *SPL*s (Figures 5F; S24). A transactivation assay demonstrated induction of the luciferase reporter coupled to 5’*SPL4* sequences by WUS, with the shortest fragment covering most of the inductive effect (Figure 5G). This sequence contains five *SPL4*^WUS^ sites, four of which could be separately tested by ChIP-PCR. This indeed confirmed direct induction of *SPL4* by WUS (Figures 5H; S25). However, these results also suggest that additional players downstream of the WUS/CLV3 feedback loop are important for the onset of flowering, which cannot be bypassed by an otherwise inductive SPL4 (Figure 5A).

### A feed-back regulation between the T6P pathway and WUS involves the age pathway

An enlarging SAM at floral transition correlates with a larger *WUS* expression domain expanding into the CZ and the outer cell layer (L1) (Figure 1D-F). Interestingly, we found reduced *TPS1* levels (*35S:amiRTPS1*) lead to an enlarged OC domain marked by *WUS* (Figure 6A) that overlaps with the CZ, marked by *CLV3* expression (Figure 6A). Plants overexpressing *TPS1* (*35S:TPS1*, Figure S2) display an increased stem cell pool with otherwise little effect on *WUS* expression (Figure 6A). The size of the CZ decreases when expressing *TPS1* under the *CLV3* promoter (Figures 6A; S2), indicating a smaller stem cell pool in support of a much smaller SAM size of the very early flowering *CLV3:TPS1* (Wahl *et al*., 2013).

*TPPJ* overexpression results in significantly enlarged SAMs (Figure 2F), and importantly, expands *WUS* expression into the outer SAM layers, while *CLV3* expression seems to be little affected (Figure 6B,C). Consistent with this, *CLV3* expression increases to much higher levels in *ML1:amiRTPPJ* (Figure 6C), suggesting an active role of the T6P pathway in the outer meristem layer regarding stem cell maintenance. In addition, *WUS* is induced (Figures 6C-D), indicating uncoupling of the WUS/CLV3 feed-back loop downstream of the T6P pathway.

We next investigate whether this is in part due to the function of the age pathway downstream of the T6P pathway. While we found *WUS* to be induced (Figure S26A,B), its expression seems to be reduced in the CZ in the transition SAM of all but *spl3,5* mutant SAMs (Figure S26A). This cannot be explained by an altered expression of *CLV3*, as we found it generally slightly reduced in all mutant lines (Figure S26C,D).

Since *WUS* and *SPL4* expression domains are largely overlapping (Daum *et al*., 2014; Olas *et al*., 2019; Torti *et al*, 2012; Yadav *et al*., 2011), we asked whether a feed-back loop, involving direct regulation of *WUS* by SPL4, would explain the observed expression patterns. We identified one *WUS^SPL4^* site −1067bp upstream of the *WUS* coding sequence (Figure 6E). SPL4 significantly and specifically reduced reporter gene expression in an Arabidopsis protoplast-based transactivation assay, indicating that it can repress *WUS in vivo* (Figure 6F,G). EMSA confirmed this result (Figure 6H). This finding is supported by a promoter study analyzing a series of deletion constructs of the 5’*WUS* sequence (Baurle & Laux, 2005). Among them one which deleted a sequence including the *WUS^SPL4^* site (Δ4, −941/-604 from transcription start site; −1067/-730 upstream of coding sequence). Importantly, this deletion construct led to a stronger activation while others rather led to a reduction of a GUS reporter, especially when affecting the sequences close to the transcription start site. This indicates the loss of regulatory sequences which are targeted by repressors of WUS such as SPL4 (Baurle & Laux, 2005). Taken together, our results demonstrate a likely transient negative feed-back regulation between SPL4 and WUS.

## Conclusion

Taken together, we unraveled dynamic feed-back regulations between sugar signaling, central meristem maintenance and flowering time regulators that coordinate cell proliferation and metabolism, providing the basis and energy necessary for doming during floral transition.

Proliferation, self-renewal and differentiation of cells originating from stem cells are the basis for the plants’ remarkable plasticity. Stem cell systems are formed during embryogenesis. The SAM as primary meristem is retained stably throughout plant development (Pfeiffer *et al*., 2017). The initiation of reproductive growth marks a very prominent change in a plant’s morphology and requires large-scale reorganization of SAM architecture and sink-source relationships within all flowering plants. Within cells, sucrose is an important basis for energy production and biomass. To prevent starvation, energy-demanding developmental transitions are tightly coordinated with endogenous sucrose availability through intricate signaling systems (Fernie *et al*, 2020). The sucrose signaling T6P pathway was previously found to impact on cell cycle regulators (Gómez *et al*., 2006). These were also demonstrated as differentially expressed at the transition stage (Klepikova *et al*., 2015).

We show that in Arabidopsis the negative feed-back loop between the central meristem maintenance regulators WUS and CLV3, considered as stable until now, uncouples during floral transition. As a result, the *WUS* expression domain is no longer fixed in the center of the SAM but stretches into the outer meristem layers demonstrating unreported OC dynamics. WUS directly regulates *TPPJ*, a component of the T6P pathway, which has an impact on SAM size and thus seems important for cell proliferation and doming of the SAM during the transition phase. Notably, organ production is stalled when *WUS* is ectopically expressed from a *CLV3* promoter into the stem cell niche, and correlates with massive proliferation of meristematic cells (Brand *et al*, 2002).

Although *TPPJ* is expressed at the rim of the PZ, it can be activated by WUS *in vivo*, requiring the cell non-autonomous function of a mobile WUS (Daum *et al*., 2014; Yadav *et al*., 2011). The fact that WUS does not activate *TPPJ* outside of this domain suggests other factors repressing it in the center of the SAM. These factors are clearly still active in a *clv3* mutant background, where the activation of *TPPJ* by WUS is restricted to the outer SAM layers, suggesting factors outside of this central regulation of the meristem maintenance network (Pfeiffer *et al*., 2017). It will be interesting to learn more about its regulation in the future.

We demonstrate that SPL4, which is expressed in a domain overlapping with the central *WUS* expression domain, directly and negatively regulates *WUS.* In turn, WUS activates *SPL4* in a likely transient negative feed-back regulation, which we suggest to balance sugar-signaling-dependent effects on meristem maintenance at transition. In addition, WUS might repress *SPL5*, as we found slightly but significantly increased expression in *wus-7* apices (Figure 5E). Although they don’t seem to respond in a WUS-dependent manner, it remains to be determined whether those SPLs expressed in the periphery of the SAM (SPL3, SPL5 and SPL9) are mobile and therefore independently report on the sucrose status to the central region of the SAM. We suggest that the combined action of the T6P pathway and SPLs expressed at the SAM determines the duration of the doming phase.

An important next step will be to understand how SnRK1 and TOR signaling (Baena-González & Lunn, 2020; Caldana *et al*, 2019; Pfeiffer *et al*., 2016) fit into the emerging picture of the mechanisms regulating floral transition in the future. Apart from the regulation of flowering time (Wahl *et al*., 2013), it is interesting that the T6P pathway also influences the re-organisation of the SAM during floral transition. Given the ubiquitous nature of carbohydrate signaling and the large-scale change in sink-source relationships within plants (Fernie *et al*., 2020), it will be interesting to determine if this regulatory mechanism is widely present in the plant kingdom.

## Materials and Methods

### Plant material and growth conditions

All Arabidopsis plants were of the Columbia accession (Col-0), except for *wus-7*, which is in the Ler background (Graf *et al*., 2010). *clv3-7* (Wisman *et al*, 1998), *clv3-10* (Forner *et al*, 2015), *35S:amiRTPS1*, *CLV3:TPS1* (Wahl *et al*., 2013), and *TCSn:GFP* (Zurcher *et al*., 2013) were described previously. The *tppj* insertion mutant (At5g65140, GABI_215C08) was obtained from the Arabidopsis resource stock center (ABRC). Growth chambers were set to 22°C in LD (16h light/8h dark) or SD (8h dark/16h light) with a light intensity of approximately 160 μmol/m^−2^s^−1^ and a relative humidity of 60-65%. Synchronized induction of flowering was performed as described (Schmid *et al*., 2003).

### Plant phenotyping

Flowering time of on average of 20 plants per genotype was scored as days to flowering, when shoots were 0.5cm (bolting), and by the total leaf number (TLN), i.e., the sum of rosette leaf (RLN) and cauline leaf numbers (CLN) (Table S1). Meristem size was measured as the area between two organ primordia under the meristem summit of a longitudinal middle section through at least 10 apices from individual plants using the Fiji software version 2.0.0-rc-69/1.52 (Schindelin *et al*, 2012).

### Generation of transgenic lines

Plants were transformed (Clough & Bent, 1998), confirmed by PCR and independent, single-insertion, homozygous T3 plants were used for all studies. Oligonucleotides for cloning and for genotyping are given in Tables S2 and S3, respectively. For *35S:TPPJ* and *35S:TPS1* lines, coding sequences of *TPPJ* (At5g65140) and *TPS1* (At1g78580) were cloned via the Gateway® entry vector *pJLBlue* reverse (Mathieu *et al*, 2007) into a *pGREEN-II*-based destination vector. Artificial microRNAs targeting *TPPJ* (*ML1:amiRTPPJ* V1 and V2, *35S:amiRTPPJ* V1 and V2) were designed with the Web MicroRNA Designer (http://wmd3.weigelworld.org/cgi-bin/webapp.cgi)(Schwab *et al*., 2006). The *Eco*RI/*Bam*HI fragment was cloned via *pJLBlue* reverse (Mathieu *et al*., 2007) into a *pGREEN-II*-based destination vector with either the *ML1* or *35S* promoter. *spl3, spl4, spl5* knockout lines were generated with the CRISPR/*Cas9* technology (Ruf *et al*, 2019; Wang *et al*, 2015). The NGG PAM recognition sites were defined using the ATUM webtool (https://www.atum.bio) (Figure S23). The *pJF1033* vector, containing a two single guided RNAs scaffold, was used as a template. *Bsa*I products were cloned into the *pJF1031* binary vector (Ruf *et al*., 2019). Plants with homozygous deletions were back-crossed to Col-0. Homozygous, *Cas9*-free lines were used to generate all higher order mutant lines.

### RT-qPCR

Total RNA was isolated by a modified phenol/chloroform method using a TRIzol reagent (phenol 38% (v/v), guanidine thiocyanate 0.8M ammonium thiocyanate 0.4M, sodium acetate 0.1M pH5.0, glycerol 5% (v/v), EDTA 5mM pH8.0, Na-lauroylsarcosine 0.5% (v/v)), followed by sodium acetate precipitation. Genomic DNA was removed with DNase I RNase-free (Ambion™/Thermo Fisher Scientific, Waltham, Massachusetts, US), and cDNA synthesis carried out using a SuperScript™IV Reverse Transcriptase Kit (Thermo Fisher Scientific, Waltham, Massachusetts, US) according to the instructions. Mature miR156 stem-loop primers (Table S4) were added to the cDNA synthesis reaction (1:1 with oligo dT(18) primer) (Varkonyi-Gasic *et al*, 2007).

RT-qPCR was performed with the ABI Prism 7900 HT fast real time PCR system (Applied Biosystems/Life Technologies, Darmstadt, Germany) using a Power SYBR^®^ Green-PCR Master Mix (Applied Biosystems/Life Technologies, Darmstadt, Germany) for expression analyses and ChIP-PCRs. Oligos are listed in Table S4 (RT-qPCR) and S5 (ChIP-PCR). For analysis the SDS 2.4 software (Applied Biosystems/Life Technologies, Darmstadt, Germany) was used (Czechowski *et al*, 2004). cDNA quality was determined and expression values were calculated (Wahl *et al*., 2013).

### RNA *in situ* hybridization

Probes were synthesized using the DIG RNA Labeling Kit (Roche, Mannheim, Germany) on CDSs of the target genes cloned into the *pGEM®-T Easy* vector (Promega, Madison, Wisconsin, US). Oligonucleotides and construct IDs are listed in Table S6. Wax embedding, sectioning and RNA *in situ* hybridization, and imaging were performed as described (Wahl *et al*., 2013).

### ChIP-PCR

For ChIP-PCR, 100 apices or 1.5g leaves per replicate (Col-0, *clv3-7* and *clv3-10*) were fixed in 1% (v/v) formaldehyde buffer (10mM sodium phosphate buffer, pH7; 50mM NaCl; 100mM sucrose) under vacuum. ChIP was performed (Kaufmann *et al*, 2010) with modifications: Antibody incubation (anti-WUS; AS11 1759; Agrisera) was extended o/n at 4°C and Agarose beads (Protein A-Agarose; sc-2001; Santa Cruz Biotechnology) to 6h at 4°C. Immunoprecipitated DNA was analyzed by RT-qPCR. Ct values of *TPPJ* and *SPL4* promoter regions were normalized to the Ct value of a region within the *UBQ10* promoter. The % of enrichment was calculated as relative to expression of the input of the individual regions. ChIP on apices of Col-0, *clv3-7* and *clv3-10* without antibody and ChIP on Col-0, *clv3-7* and *clv3-10* leaves were controls (Figure S10). Note that in contrast to the input no amplification was observed in the controls (Figure S11, S25) and the previously published confirmed WUS target sites were detected (Figure S12)(Leibfried *et al*., 2005; Yadav *et al*., 2011). Oligonucleotides are listed in Table S5.

### Transactivation assay

For the effector line, the *WUS* (At2g17950) and *SPL4* (At1g53160) CDS were cloned via the Gateway® entry vector *pJLBlue* reverse (Mathieu *et al*., 2007) into the *pMDC32* Gateway® vector (pMML058) and the *pGW5* (pKK77). For the reporter constructs, designated *5’TPPJ* regions were *Kpn*I/*Acy*I cloned into the Gateway® *pMDC162* vector to obtain GUS fusions and *5’SPL4* and *5’WUS* regions were *Kpn*I/*Spe*I cloned into *pGREEN800II-LUC* (Hellens *et al*, 2005)(Table S2). *35Somega:LUC-NOS* was used as transformation control in the *5’TPPJ* assays. Protoplasts were isolated from 4-week-old plants, transfected (Wu *et al*, 2009) and incubated for 20h at 22°C and 100µmol/m^−2^s^−1^. Luciferase activity of the *5’TPPJ, 5’SPL4* and *5’WUS* assays was measured with a luciferase and dual-luciferase reporter assay system, respectively (Promega, Madison, Wisconsin, US). GUS activity was determined as described (Yoo *et al*, 2007).

### EMSA

The *WUS* CDS without STOP codon was cloned via the Gateway® entry vector *pDONR207* (pMML059) into a Gateway® destination vector (*pDEST24*). *35S:WUS-GST* (pMML063) was transformed into *Escherichia coli* Rosetta plysS cells and protein production induced with 1mM isopropyl β-D-1-thiogalactopyranoside at 30°C o/n. Cell lysis was performed with 1x sonication of 5 sec, 20% power, 4 cycles (Sonoplus Hd 2070 Sonicator, Badelin, Berlin, Germany) in a freshly prepared buffer (20mM Na-phosphate buffer, pH7.4; 0.5MNaCl; 1mM phenylmethylsulfonyl fluoride; 1mM Ethylenediaminetetraacetic acid; 1 tablet cOmplete™ Protease Inhibitor Cocktail (Merck, Darmstadt, Germany) per 10ml buffer). 2µg protein (crude extract, determined by Pierce^TM^ BCA Protein Assay Kit, Thermo Fisher Scientific, Waltham, Massachusetts, US) was used for the EMSA. 50bp, double-stranded, 5′-IRDey-682-labeled probes, spanning the putative *TPPJ^WUS^* and *WUS^SPL4^* sites are listed in Table S7 (5’*TPPJ*) and Table S8 (5’*WUS*). Binding reactions were carried out with the Odyssey® EMSA Kit (LI-COR^®^) according to the instructions with a competitor and/or mutated competitor to probe ratio of 1:200 and visualized with the Oddysey Infrafed Imaging System (Li-Cor, Lincoln, NE).

### Confocal microscopy

SAMs of *a TCSn:GFP* cytokinin reporter line (Zurcher *et al*., 2013) were imaged with a Leica SP8 confocal laser scanning system equipped with a M6000B-CS microscopy stage, an Argon laser (65mV), and a 40x water immersion HCX APO objective as follows: Laser output power 20%; GFP excitation (green, wavelength 488nm), emission detection channel 3 (495-520nm), gain PMT 800V; plastid auto-fluorescence (blue), emission detection channel 4 (700-800nm), gain PMT ∼500V; scan speed 600Hz in xyz bi-directional scanning mode with a z-stack distance of approx. 10µm. Offset=0; pixel dimension: 1024×1024. Middle sections of representative SAMs were extracted from z-stacks using the Fiji software version 2.0.0-rc-69/1.52 (Schindelin *et al*., 2012).

### Statistical consideration

Statistical significance was analyzed by both one-way ANOVA (Analysis of Variance) with Tukey’ Post Hoc HSD (Honestly Significant Difference) based on Tukey-Kramer correction (P value <0.05) or a two-tailed Student’s *t*-test.

## Acknowledgments

The authors wish to thank all members of the Wahl group for fruitful discussions, M. Schmid and P. Wigge for critical reading of the manuscript, D. Walter for input on statistical analyses, K. Kontbay for providing the SPL4 effector construct, M. Liang and J. Van Dingenen for their support with the *spl* CRISPR/*Cas9* deletion lines, M. Molochko for sharing the *35Somega:LUC-NOS* plasmid, J. Forner for providing the *pJF1031* and *pJF1033* plasmids. Work in the Wahl group was supported by the BMBF (SolaMI, 031B0191), the DFG (SPP1530: WA3639/1-2), and the Max Planck Society.

## Author Contributions

VW conceived and designed the experiments and prepared the figures with contributions from MML. VW and MML analyzed the data and wrote the manuscript. MML, KF, AK, and VW performed all essential experiments, i.e. VW generated most of the lines used, except for *ML1:amiRTPPJ* (KF), the *spl* CRISPR/Cas9 deletion lines (MML), and all crosses thereof (MML). MML performed all transactivation assays, ChIP-PCRs, EMSAs and RT-qPCR analyses. MML, AK, KF, CA, and VW analyzed SAM sizes and scored flowering time. RNA *in situ* hybridizations were performed by VW, AK, KF and MML. FK and VW took the confocal images. All authors have read and commented on the text and figures within this manuscript.

## Data Availability Section

Supporting information is provided in a separate file. This includes:

Figures S1 to S26

Tables S1 to S8

SI References

## Supplementary Figures

**Fig. S1.**
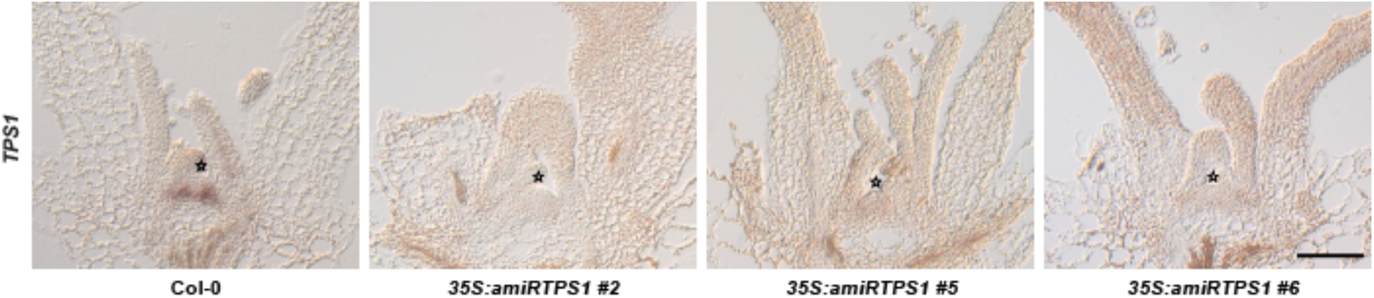
*TPS1* expression at the shoot apical meristem of *35S:amiRTPS1* plants. Downregulation of *TPS1* by overexpressing an artificial microRNA targeting *TPS1* (*35S:amiRTPS1*) in plants of three independent lines (#2, #5 and #6; (Wahl *et al*, 2013)) as analyzed by RNA *in situ* hybridization using a specific probe against *TPS1*. Pictures depict representative longitudinal middle sections through vegetative apices (8 DAG in LD) of Col-0 and *35S:amiRTPS1*, respectively. Line #6 was used for the experiments in this study. Star indicates SAM summit. Scale bar is 50 µm.

**Fig. S2.**
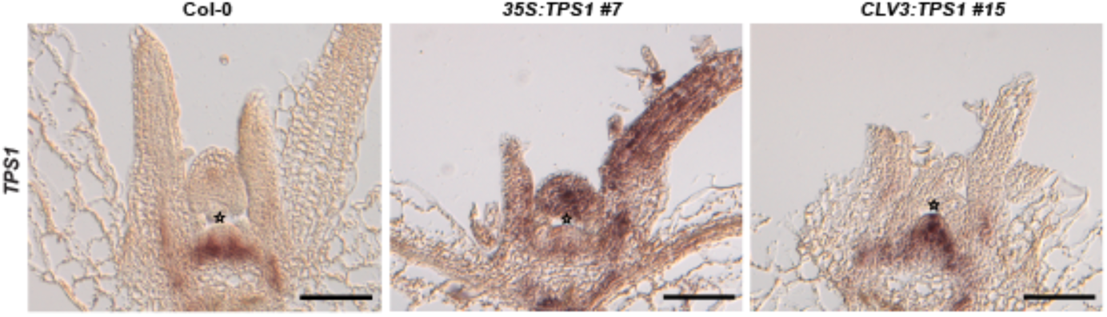
*TPS1* expression at the shoot apical meristem of *35S:TPS1* and *CLV3:TPS1* plants. In comparison to expression in Col-0 *TPS1* was generally found upregulated in plants overexpressing *TPS1* (*35S:TPS1*, #7) and the central zone when expressing it under the control of the *CLAVATA3* (*CLV3*) promoter (*CLV3:TPS1*, #15) as analyzed by RNA *in situ* hybridization using a specific probe against *TPS1*. Pictures depict representative longitudinal middle sections through vegetative apices (8 DAG in LD) of Col-0, *35S:TPS1*, and *CLV3:TPS1*, respectively. Star indicates SAM summit. Scale bars are 50 µm.

**Fig. S3.**
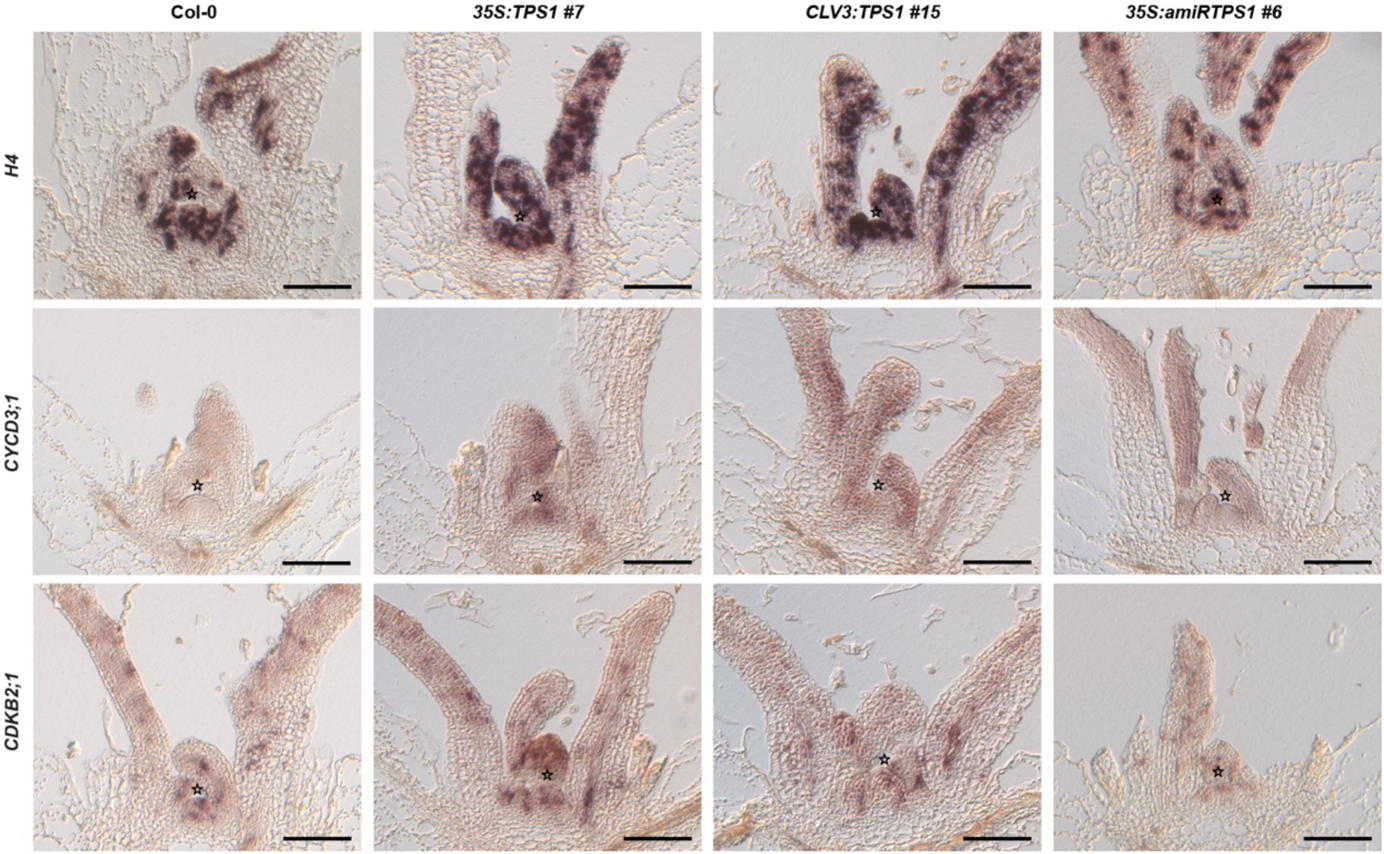
Cell cycle gene expression at the shoot apical meristem of *35S:TPS1*, *CLV3:TPS1* and *35S:amiRTPS1* plants. RNA *in situ* hybridization using specific probes for *HISTONE H4* (*H4*), *CYCLIN D3;1* (*CYCD3;1*) and *CYCLIN-DEPENDENT KINASE 2;1* (*CDKB2;1*) to illustrate expression in longitudinal middle sections through the SAM of Col-0, *35S:TPS1* (#7), *CLV3:TPS1* (#15) and *35S:amiRTPS1* (#6). Star indicates SAM summit. Scale bars are 50 µm.

**Fig. S4.**
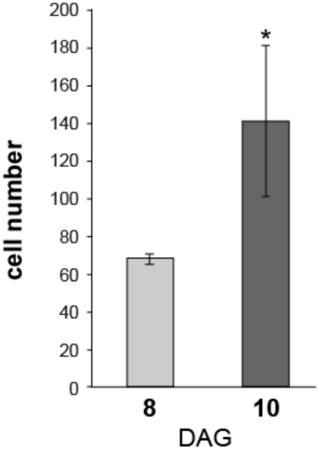
Metrics at the SAM before and at the floral transition. Cell numbers as recorded at 8 (vegetative SAM) and 10 days (transition SAM) after germination (DAG). Significance calculated based on one-way Anova; *P<0.05.

**Fig. S5.**
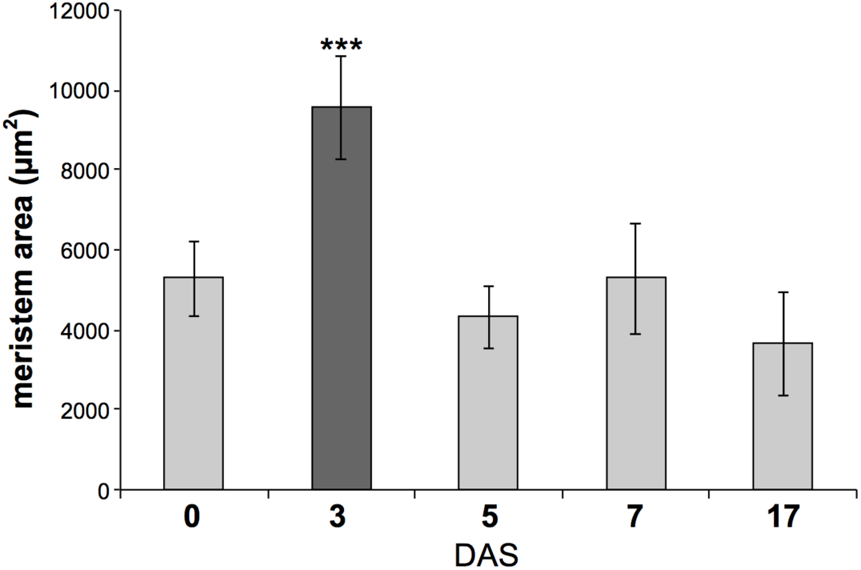
Shoot apical meristem area of SD-grown and LD-induced plants. Vegetative and inflorescence SAM areas (light grey), and transition SAM area (dark grey) of wild-type plants grown in SD for 30 days and transferred to LD for 3, 5, 7 and 17 days (DAS). Error bars denote s.d.; \*\*\**P*<0.001 in relation to 0 days after transfer to LD (Student’s t-test).

**Fig. S6.**
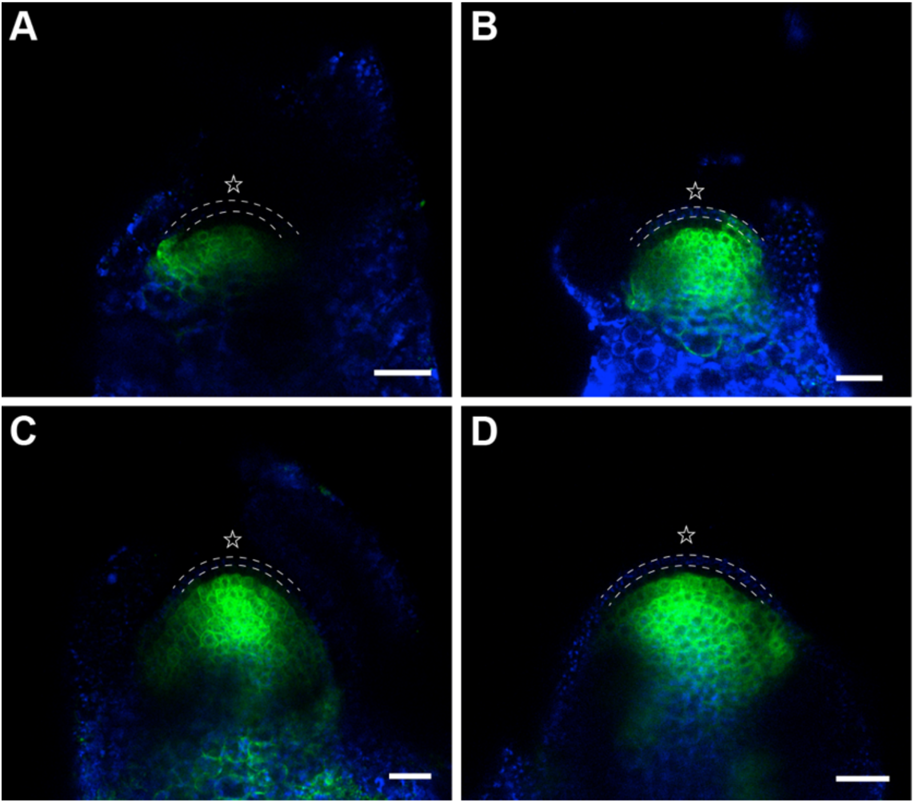
Analysis of the cytokinin signaling reporter TCSn:GFP. Representative confocal laser scanning microscope images of SAMs of (*A*) 6 DAG, (*B*) 8 DAG, (*C*) 10 DAG, and (*D*) 12 DAG of TCSn:GFP cytokinin reporter line (green (Zurcher *et al*, 2013); blue – autofluorescence of chloroplasts). Please note the absence of cytokinin signal in the L1 (between dashed lines) and L2 (cell layers below) in the meristem proper. Star indicates SAM summit, scale bars are 25μm.

**Fig. S7.**
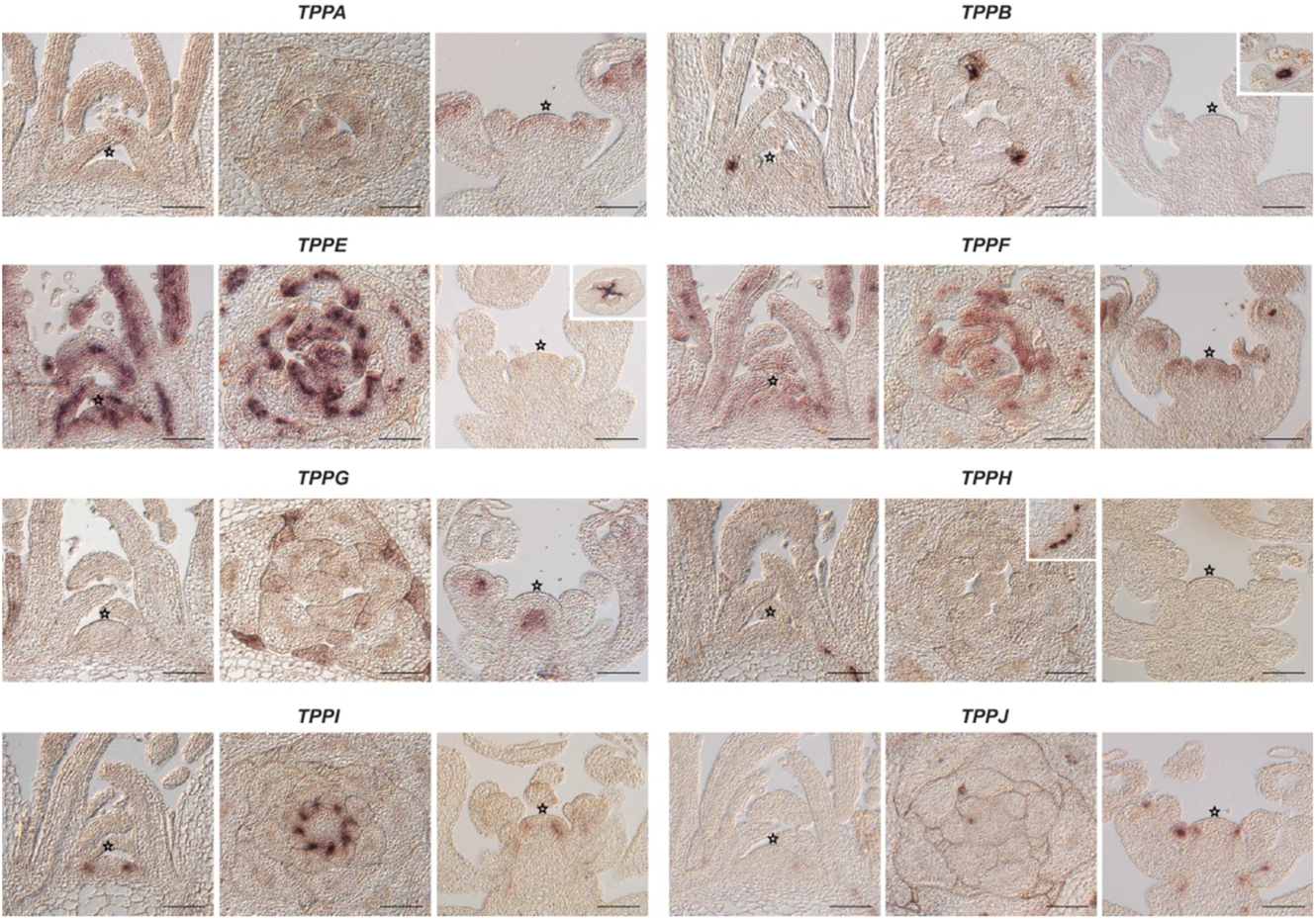
Expression of Arabidopsis *TPP* genes at the shoot apical meristem detected by RNA *in situ* hybridization. The individual panels contain: a longitudinal middle section and a cross section through a vegetative Col-0 apex and a longitudinal middle section through an inflorescence apex. Representative pictures of *TPPC* and *TPPD* were omitted from this figure, since their specific probes gave no signal, indicating that transcripts of *TPPC* and *TPPD* were below detection limit in the tissues and/or stages analyzed. Star indicates SAM summit. Scale bars are 100 μm.

**Fig. S8.**
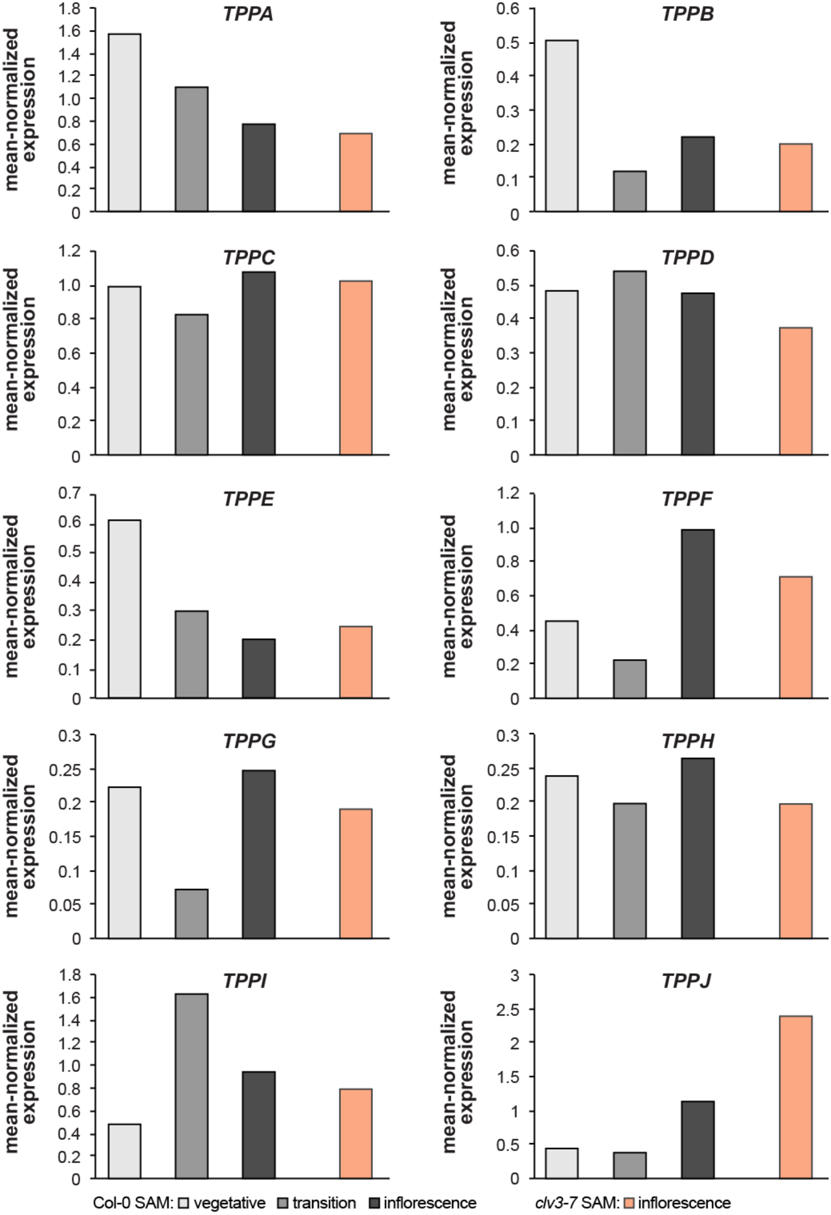
AtGenExpress expression profiles of the Arabidopsis *TPP* genes. Values for Col-0 vegetative, transition and inflorescence apices (grey scale) and *clv3-7* inflorescence apices (orange) extracted from data deposited for (Schmid, *et al*, 2005). All data points were mean-normalized.

**Fig. S9.**
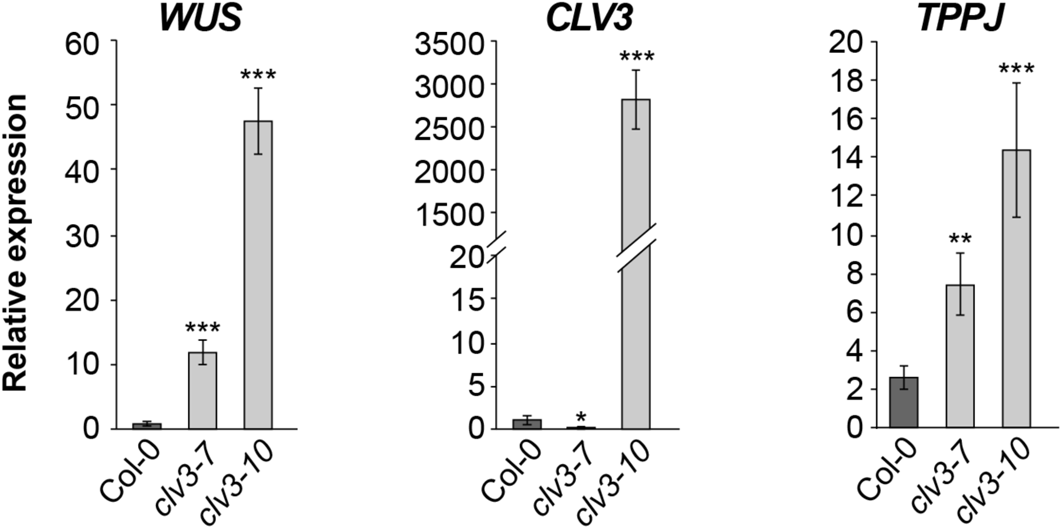
Expression of meristem maintenance genes and *TPPJ* in the *clv3-7* and *clv3-10* mutant SAM. RT-qPCR results of *TPPJ*, *WUS* and *CLV3* expression in SD-grown *clv3-7* and *clv3-10* SAMs, harvested 40 days after germination. Please note that the graph for *TPPJ* is a biological repetition of the graph in Figure 2C. Please also note that *clv3-10* is a TALEN generated mutant with a five nucleotide deletion in the last exon, which renders the transcript inactive to produce a functional protein (Forner *et al*, 2015). This is the cause for increased expression of *CLV3* downstream of increased WUS in the *clv3-10* mutant background. Error bars denote s.d.; \**P*<0.05, \*\**P*<0.01, \*\*\**P*<0.001 (one-way ANOVA).

**Fig. S10.**
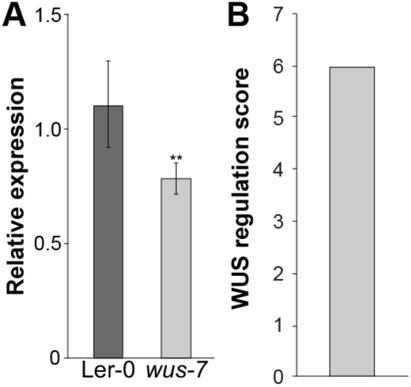
*TPPJ* Expression in *WUS* transgenic lines. (*A*) RT-qPCR results of *TPPJ* expression in SD-grown *wus-7* as compared to wild-type Ler-0 SAMs, harvested 30 days after germination. Error bars denote s.d.; **P<0.01, (one-way ANOVA). (*B*) *TPPJ* expression is increased upon WUS induction as shown by a WUS regulation score (WRS). According to Busch and colleagues, the WRS quantifies gene expression in response to a modulation in WUS activity normalized to the average response of all genes. Values extracted form a data set, deposited as a supplemental material of a publication by (Busch *et al*, 2010).

**Fig. S11.**
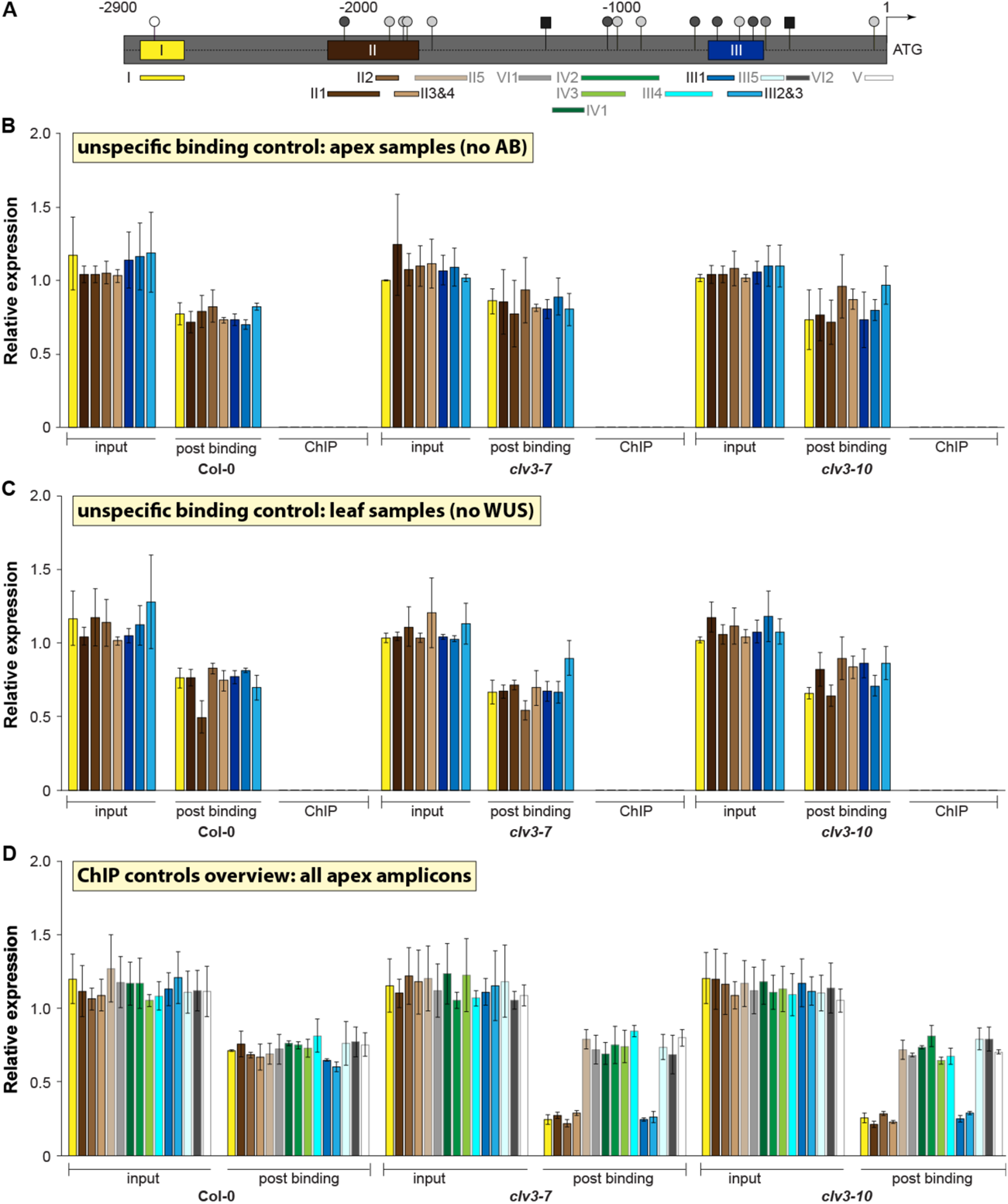
ChIP-PCR control experiments for Figure 3C-E. (*A*) Overview of *5’TPPJ* intergenic sequence with all investigated putative *TPPJ^WUS^* sites (grey circles, black boxes), position of ChIP-PCR amplicons corresponding to the results shown in (B-D). Black framed boxes marked with I, II (1-4), and III (1-3) indicate *5’ TPPJ* regions directly bound by WUS – I: −2795 – −2789 bp, II: −2073 – −1830 bp, and III: −652 – −564 bp, as presented in the Figure 3. Grey framed boxes (II5, III3-4, IV1-3, V1, VI1-2) represent *5’TPPJ* regions not directly bound by WUS. (*B-D*) Relative expression of investigated regions (I-III) containing *TPPJ^WUS^* elements (I, II1-4, III1-2) in Col-0, *clv3-7* and *clv3-10* (*B*) shoot apex samples without WUS antibody, and (*C*) leaf samples without WUS were used as specificity controls. Please note the amplification in the input and post-binding fractions (PB) and the absence of amplification in the ChIP samples indicating specificity of the antibody used. (*D*) Control overview of all investigated putative *TPPJ^WUS^* sites in input and post binding fraction samples. Error bars denote s.d.

**Fig. S12.**
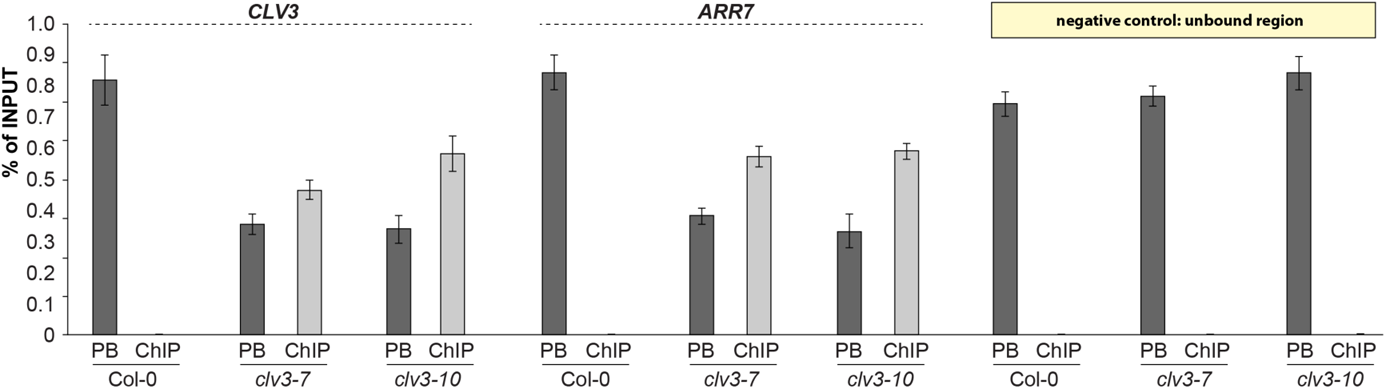
ChIP-PCR on known WUS targets using a specific WUS antibody. Relative to the input enrichment of known *CLV3^WUS^* (Yadav *et al*, 2011) and *ARABIDOPSIS RESPONSE REGULATOR 7* (*ARR7^WUS^*) (Leibfried *et al*, 2005) in *clv3-7* and *clv3-10* as measured by ChIP-PCR. As a negative control a *5’TPPJ* region was used that did not show any WUS binding (unbound region). Error bars denote s.d.

**Fig. S13.**
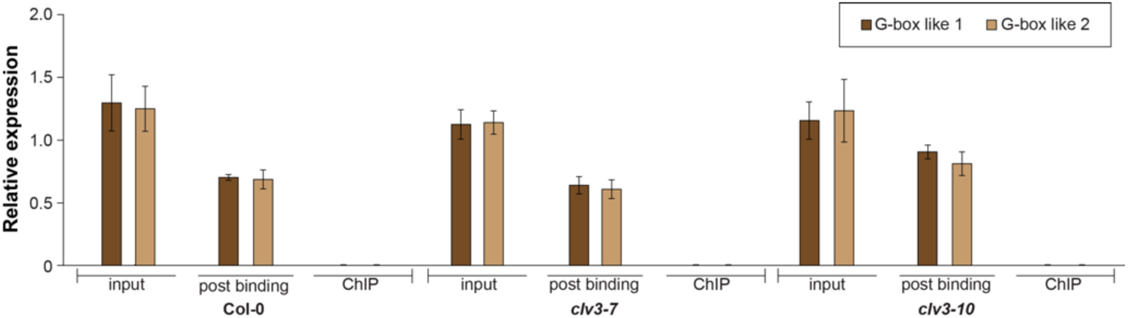
ChIP-PCR on G-box like WUS binding sites. Relative expression of two G-box like WUS binding regions (CACGTG) (Sloan *et al*., 2020) present in the *5’TPPJ* intergenic region in input, post binding and ChIP samples of Col-0, *clv3-7* and *clv3-10*. G-box like 1 and G-box like 2 correspond to the regions VI1 and VI2 of Fig. S11A. Please note the amplification in the input and post-binding fractions and the absence of amplification in the ChIP samples. Error bars denote s.d.

**Fig. S14.**
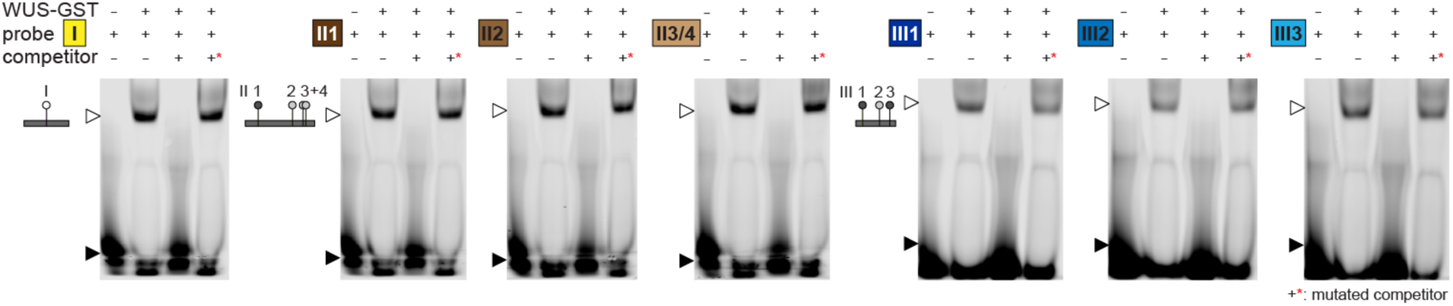
EMSA control experiments. EMSA for WUS binding to the indicated regions (I-III) in Figure 3A as a control for Figure 3F. Shifted band in the presence of WUS protein (open arrow head), non-shifted fraction (closed arrow head). Please note the shifted band in the presence of the mutated competitor (x*, Table S8) indicating specificity of the binding.

**Fig. S15.**
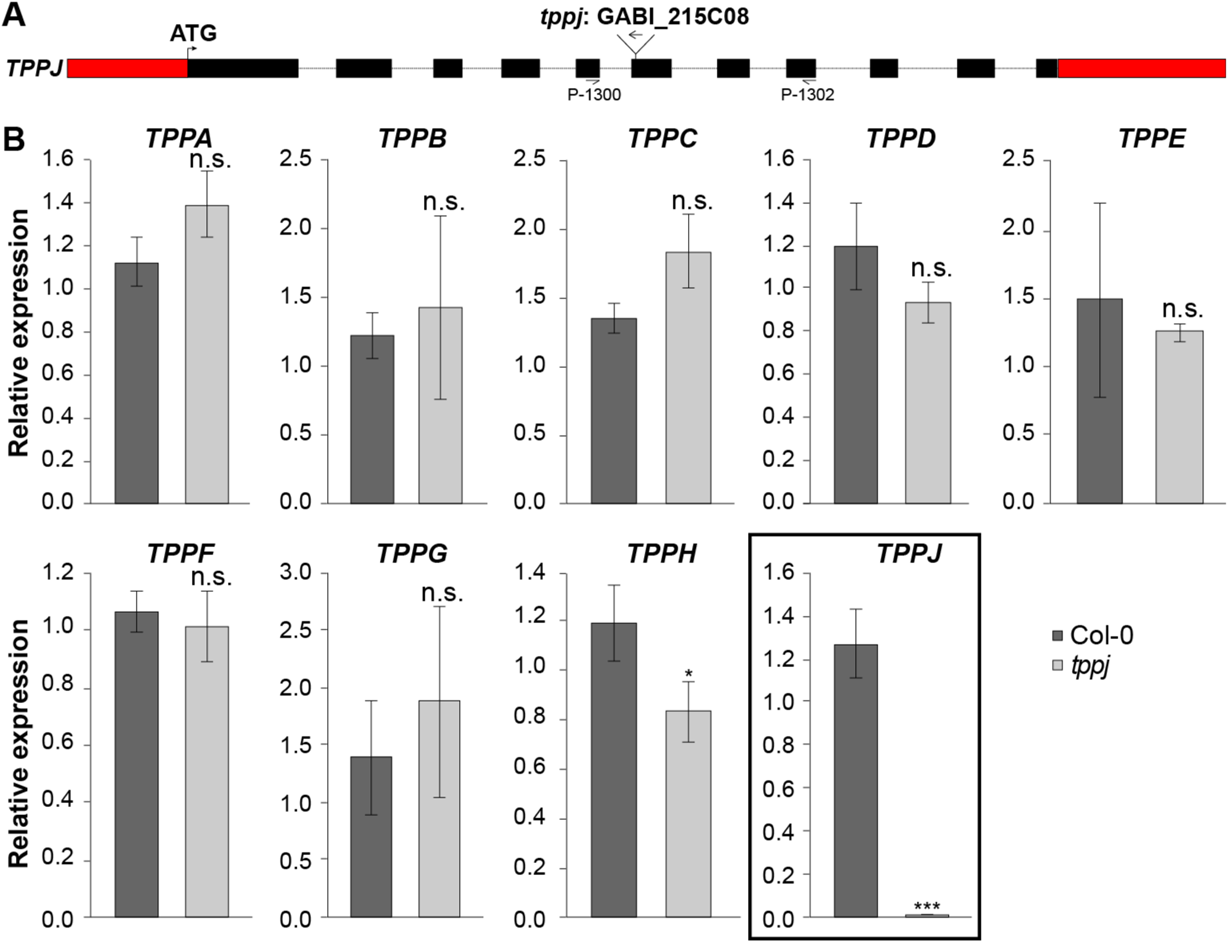
Chracterization of the GABI_215C08 line. *(A)* Schematic illustration of the T-DNA insertion in the 6^th^ exon of *TPPJ* including positions of oligos used for genotyping. *(B)* Expression of *TPPs* in LD-grown Col-0 and *tppj* rosettes, harvested without roots at 10 days after germination as measured by RT-qPCR. In this experiment *TPPI* was not detectable. Error bars denote s.d.; *<0.01; ***P<0.001 (one-way ANOVA).

**Fig. S16.**
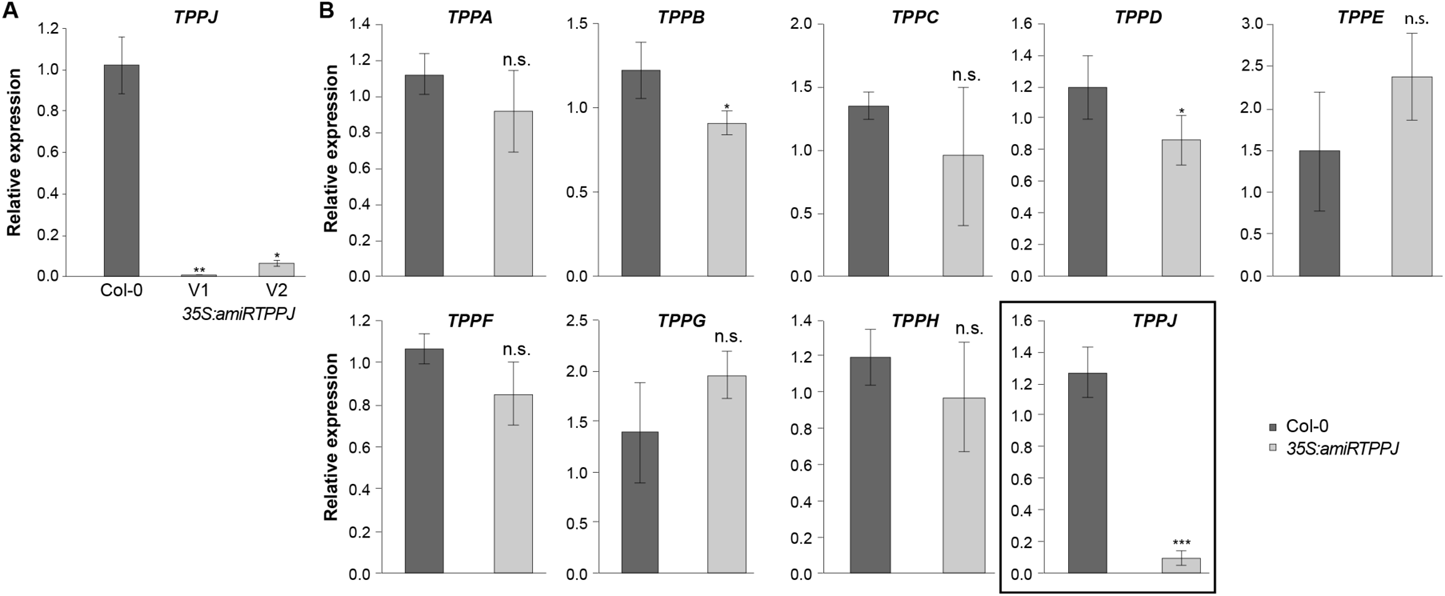
Characterization of the *35S:amiRTPPJ* lines. *(A)* Downregulation of *TPPJ* expression in *35S:amiRTPPJ* V1 and V2 rosettes shown by RT-qPCR, relative to Col-0. *(B)* Expression of all *TPPs* in LD-grown Col-0 and *35S:amiRTPPJ* V1 rosettes, harvested without roots at 10 days after germination by RT-qPCR. In this experiment *TPPI* was not detectable. Error bars denote s.d.; \*\**P*<0.01 (one-way ANOVA).

**Fig. S17.**
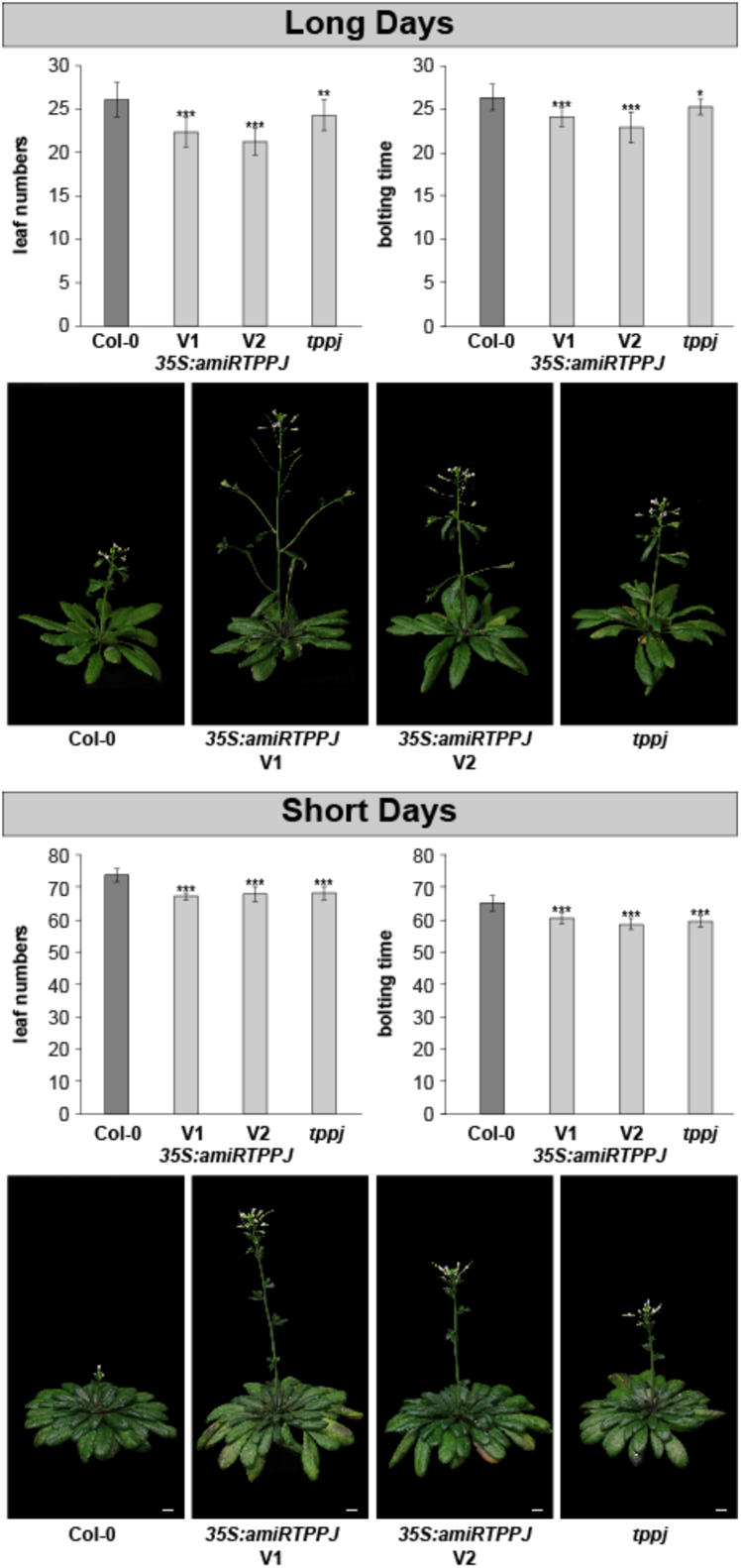
Earlier flowering of *35S:amiRTPPJ* and *tppj* lines. Flowering time of *35S:amiRTPPJ* V1 and V2 and *tppj* T-DNA insertion line as shown by the total leaf numbers and days to bolting, relative to the wild type in LD and in SD. V1 and V2 indicate two independent versions of artificial microRNAs designed to target *TPPJ* transcript. Representative pictures of *35S:amiRTPPJ* V1 line with stronger reduction in *TPPJ* expression, *35S:amiRTPPJ V2* and *tppj* in comparison to Col-0 are depicted. Scale bars are 1 cm. Error bars denote s.d.; significance calculated based on one way Student’s t-test; *P<0.05, **P<0.01, ***P<0.001.

**Fig. S18.**
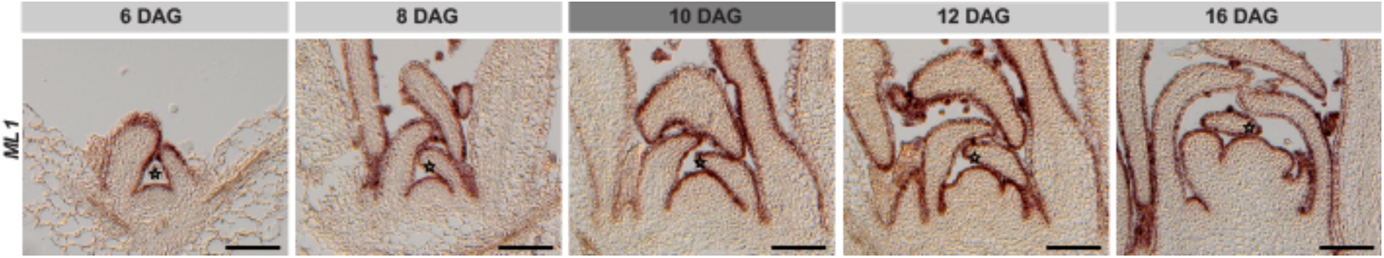
*ML1* expression throughout development. RNA *in situ* hybridization using a specific probe against *ML1* on longitudinal sections through Col-0 apices from plants grown in LD. 6, 8 DAG depict vegetative SAMs, 10 DAG the transition to flowering, and 12, 16 DAG inflorescence SAMs. Star indicate SAM summit. Scale bars are 100 μm.

**Fig. S19.**
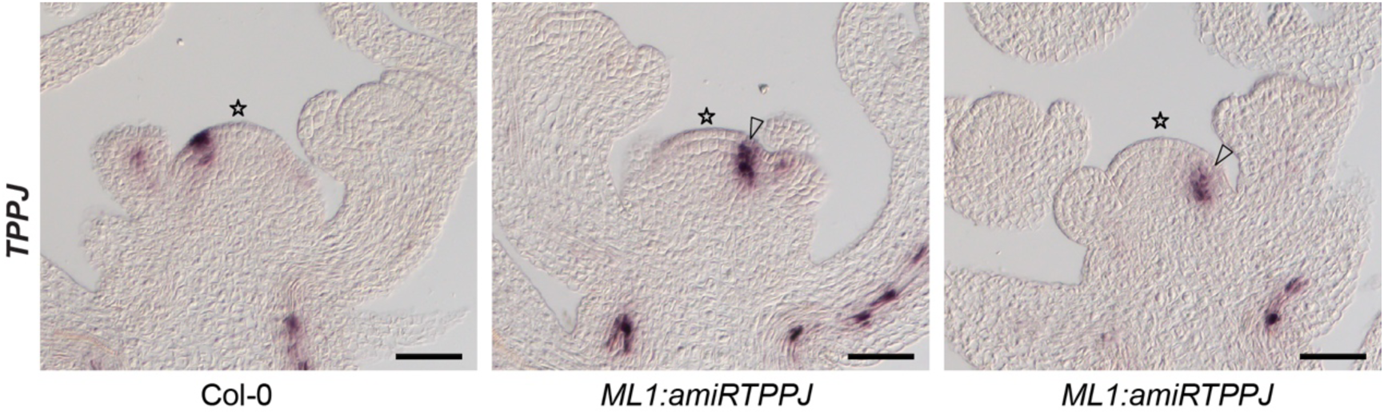
*TPPJ* expression at the SAM in the background of *ML1:amiRTPPJ* plants. RNA *in situ* hybridization with a specific probe against *TPPJ* did not detect *TPPJ* transcript in L1 (arrow head) on longitudinal sections through SAMs of *ML1:amiRTPPJ* plants (V1). Pictures depict two biological replicates of representative inflorescence SAMs of *ML1:amiRTPPJ* in comparison to Col-0. Star indicate SAM summit. Scale bars are 50 µm.

**Fig. S20.**
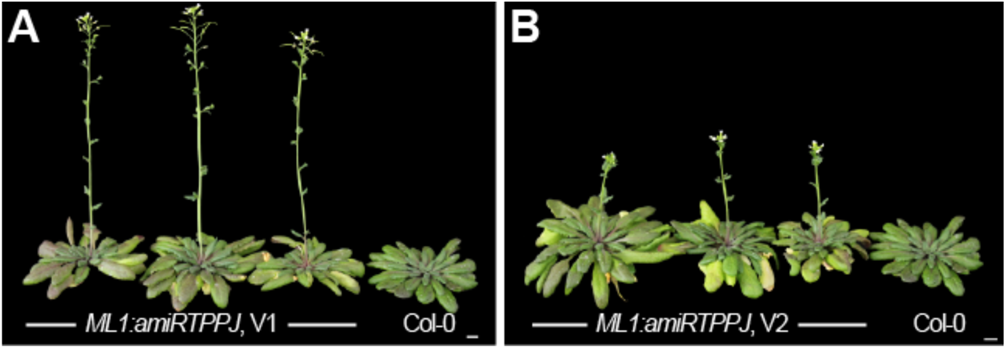
Early flowering phenotype of *ML1:amiRTPPJ* lines. Representative pictures of *(A)* the stronger *ML1:amiRTPPJ* V1 line and *(B) ML1:amiRTPPJ* V2 in comparison to Col-0 grown in SD conditions. Scale bars are 1 cm.

**Fig. S21.**
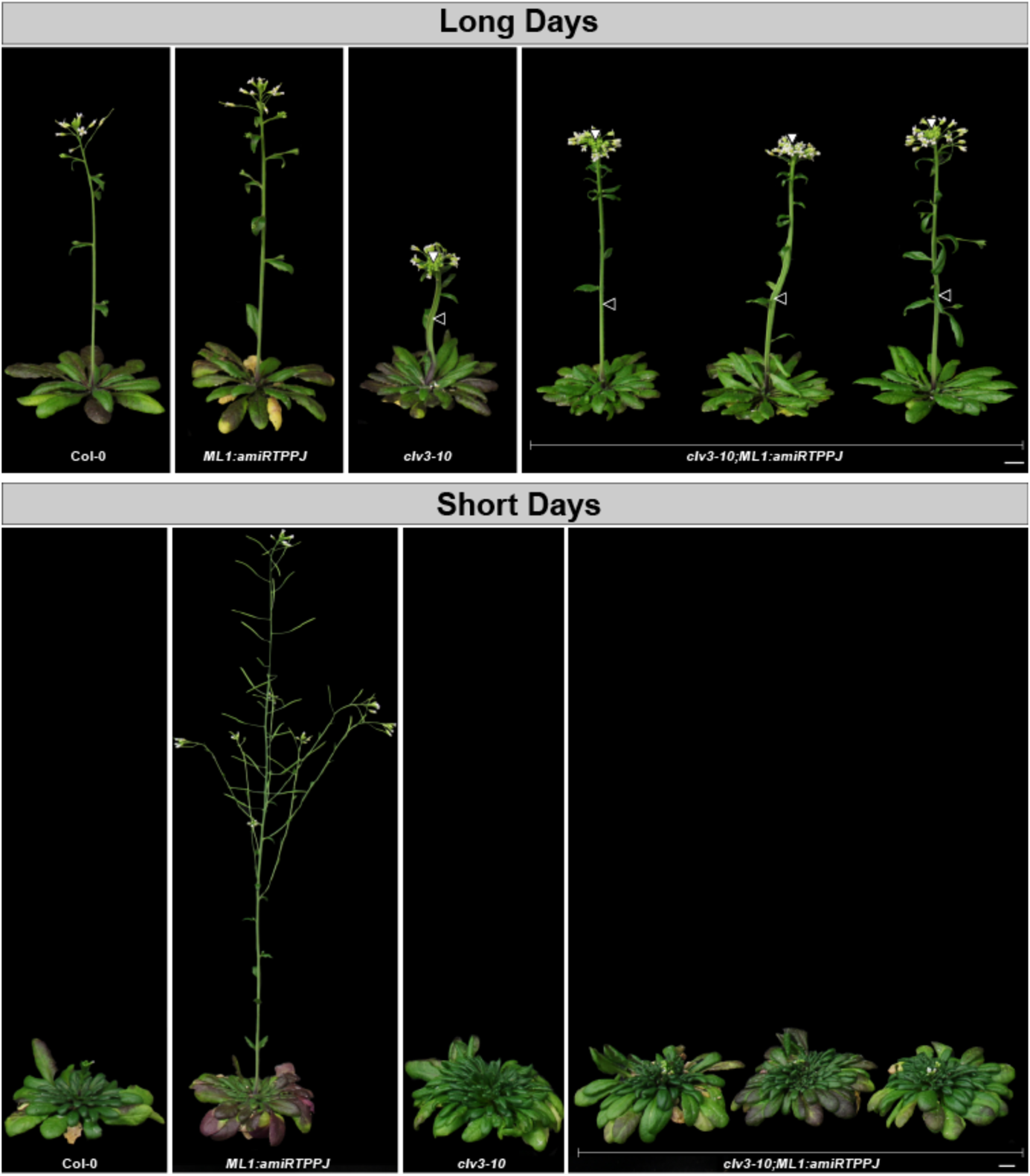
Wild-type like flowering phenotype when *ML1:amiRTPPJ* is introgressed into *clv3-10*. Representative pictures of *ML1:amiRTPPJ V1;clv3-10* next to the parental lines and wild type (Col-0) in LD (upper panel) and SD (lower panel) indicating a wild-type like flowering phenotype. Open and closed arrowheads indicate fasciated stem and enormous SAM, respectively. Note the reduced phenotype of the crosses as compared to the *clv3-10* mutant. Scale bars are 1 cm.

**Fig. S22.**
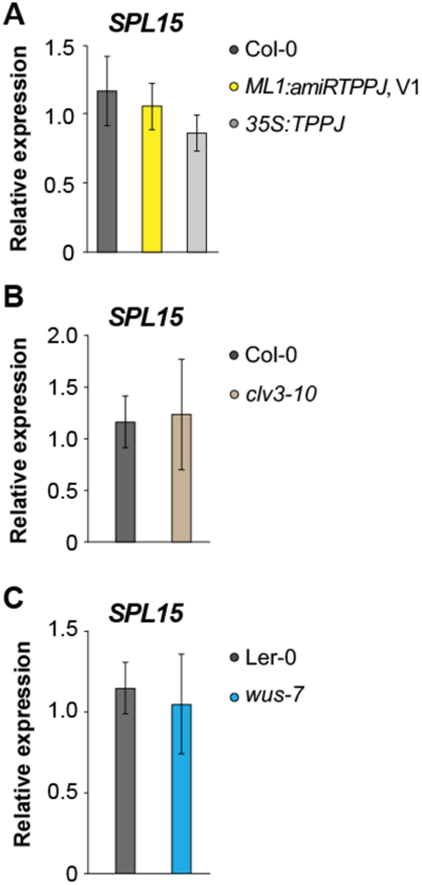
Expression of *SPL15* at the SAM. *SPL15* does not significantly change at the SAM of *(A) ML1:amiRTPPJ* and *35S:TPPJ*, *(B) clv3-10*, and *(C) wus-7* compared to its expression in the corresponding wild type. Statistical significance was calculated using one-way ANOVA. Error bars denote s.d.

**Fig. S23.**
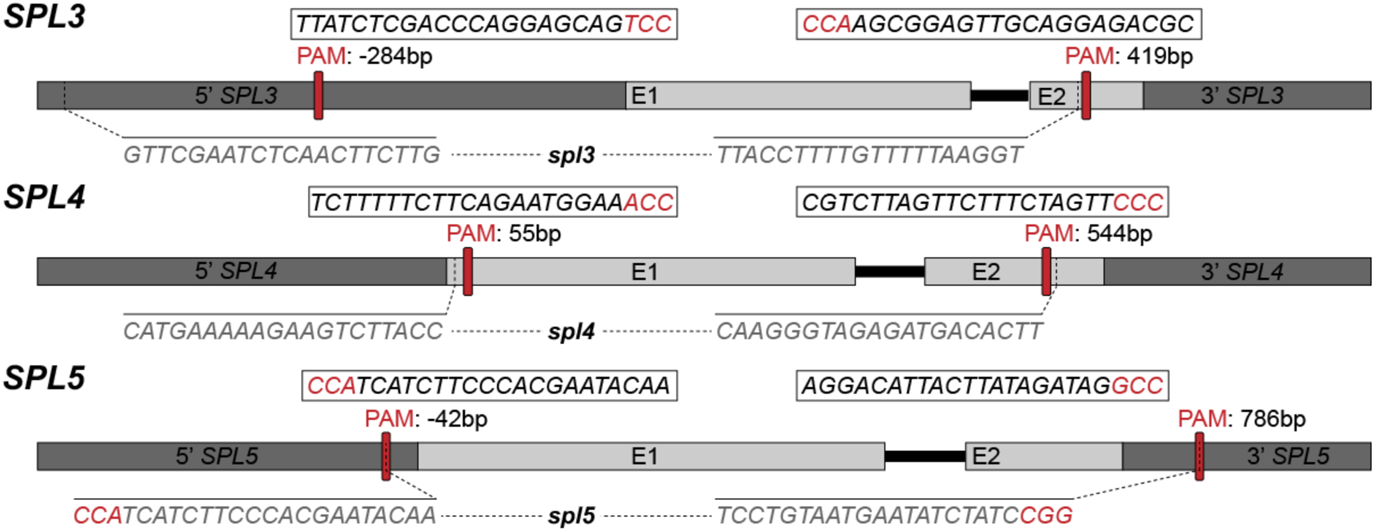
*SPL3*, *SPL4*, and *SPL5* mutant lines. Schematic illustration of the deletions in the *SPL3* (At2g33810), *SPL4* (At1g53160), and *SPL5* (At3g15270) locus generated by CRISPR/Cas9. Positions of the PAM sequence are given relative to the transcription start site (ATG). Sequences of guided RNAs with PAM sites (in red) are given in white boxes. In grey the sequences flanking the deletion are presented. E – exon.

**Fig. S24.**
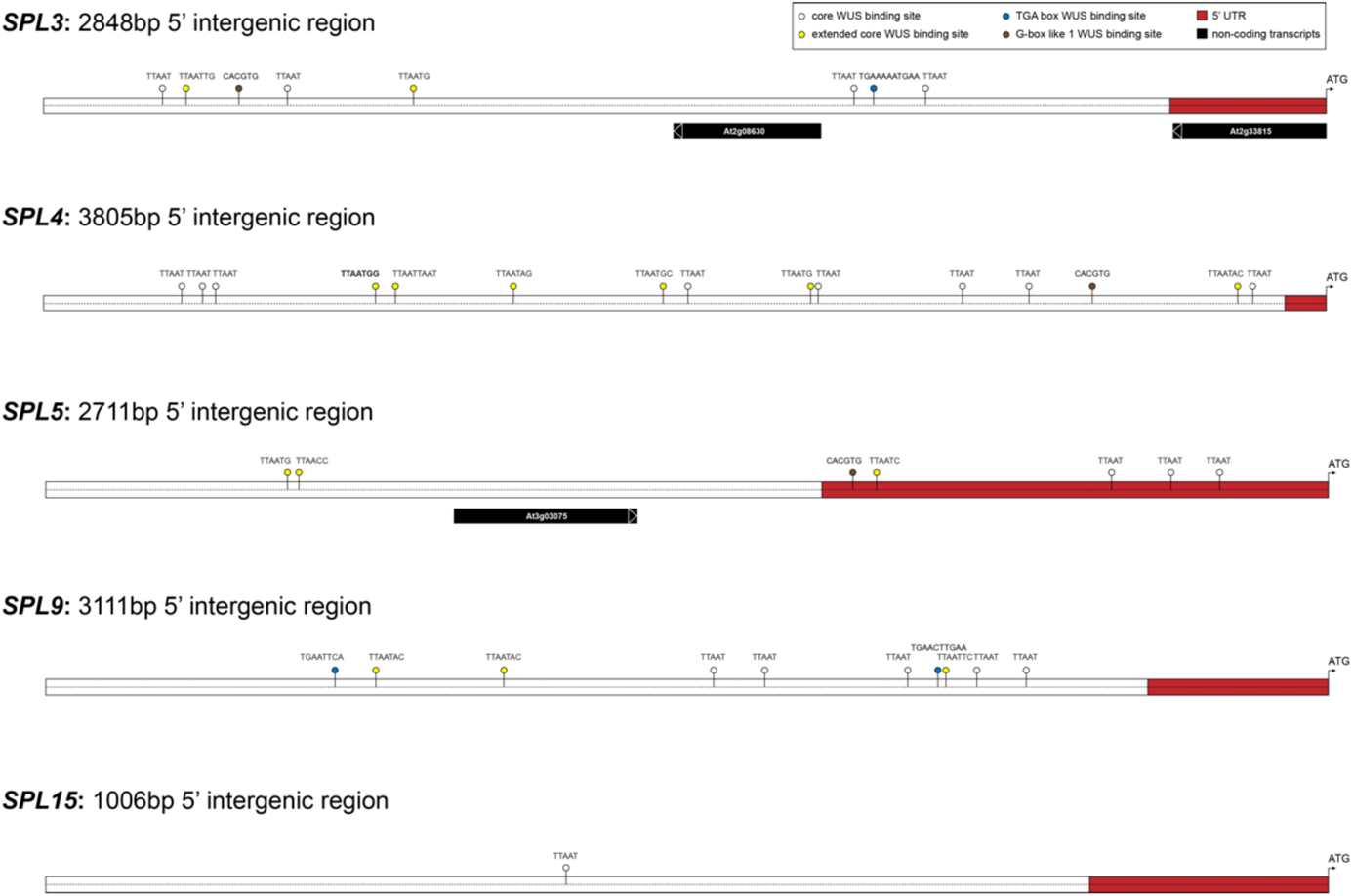
Potential *SPL^WUS^* sites in the 5’ intergenic regions of *SPL3*, *SPL4*, *SPL5*, *SPL9*, and *SPL15*. Overview of analyses of *SPL3*, *SPL4*, *SPL5*, *SPL9* and *SPL15* 5’ intergenic sequences for the presence of putative core binding sites (white circles), extended binding sites (yellow circles), TGA boxes (green circles) and G-box-like sites (brown circles). Red boxes indicate *5’UTRs*. Black boxes represent putative transcripts annotated in the intergenic regions upstream the ATG of *SPL3* and *SPL5*. The total sequence lengths are given in base pairs (bp).

**Fig. S25.**
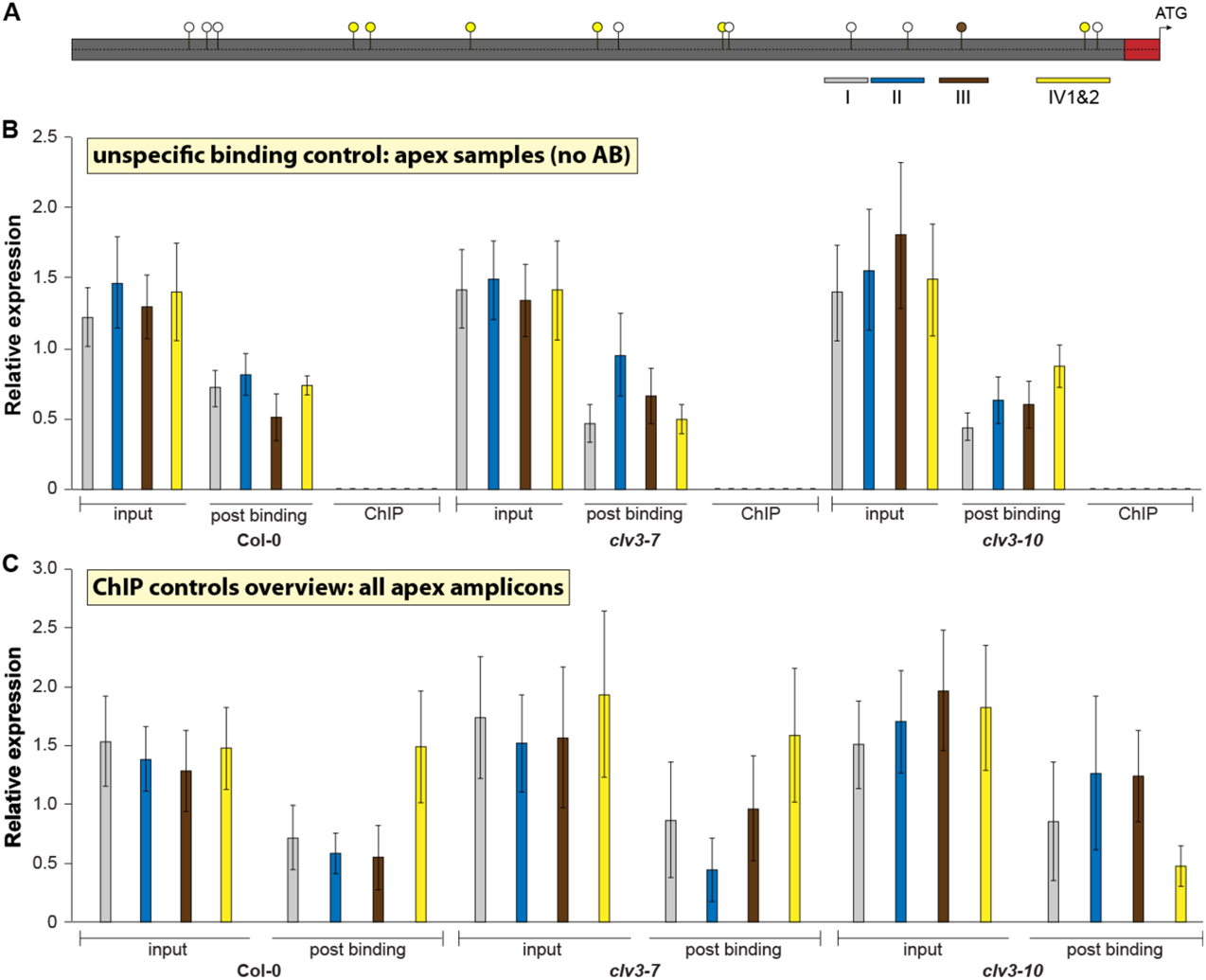
ChIP-PCR control experiments. (*A*) Overview of 5’ *SPL4* intergenic sequence with all investigated putative *SPL4^WUS^* sites (white, yellow and brown circles), position of ChIP-PCR amplicons corresponding to the results shown in (B and C). Black framed boxes marked with I, II, III, and VI (1&2) indicate *5’SPL4* regions directly bound by WUS – I: −1073 – −1068 bp, II: −880 – −875 bp, III: −697 – −691 bp, and IV: −259 – −209, as presented in Figure 5H. (*B*,*C*) Relative expression of investigated regions (I-IV) containing *SPL4^WUS^* elements in (*B*) Col-0, *clv3-7* and *clv3-10* shoot apex samples without WUS antibody. (*C*) Control overview of all investigated putative *SPL4^WUS^* sites in input and post binding fraction samples. Error bars denote s.d.

**Fig. S26.**
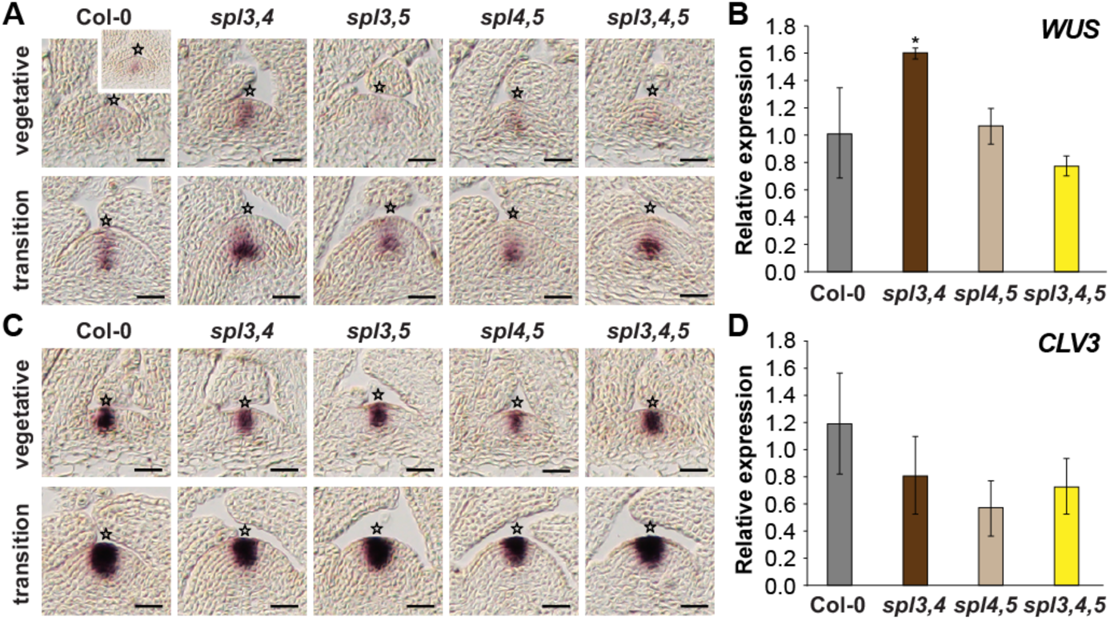
*WUS* and *CLV3* expression in apices of SD-grown SPL double and triple CRISPR/*Cas9* deletion mutants. Expression provided as *(A,C)* RNA *in situ* hybridization and *(B,D)* RT-qPCR. *WUS (A,B)* and *CLV3* (C,D). n=3. Error bars denote s.d.; significance calculated based on one-way ANOVA; \**P*<0.05. Scale bars: 25µm. Star indicates SAM summit.

## Supplementary Tables S1 to S7

**Table S1.** Flowering times of Col-0, mutants and transgenic plants. DTF, days to flowering (equals bolting time, see before); RLN, rosette leaf number; CLN, cauline leaf number; TLN, total leaf number; n, number of individual plants; p-value of student’s *t*-test for DTF with Co-0; LD, long days; SD, short days; NA, not analyzed (leaf number above 200).

|  | DTF | DTF range | RLN | CLN | TLN | TLN range | n | p-value |
| --- | --- | --- | --- | --- | --- | --- | --- | --- |
| **Presented in Figure 4 of the main text:** |  |  |  |  |  |  |  |  |
| **Experiment 1 (SD)** |  |  |  |  |  |  |  |  |
| Col-0 | 71.6 | 69-75 | 66.4 | 11.5 | 77.9 | 76-82 | 20 | - |
| *ML1:amiRTPPJ V1 #1* | 55.0 | 50-58 | 49.4 | 12.0 | 61.3 | 58-63 | 20 | 3.29E-22 |
| *ML1:amiRTPPJ V2 #5* | 64.4 | 61-66 | 52.8 | 9.2 | 62.0 | 60-64 | 20 | 3.07E-14 |
| **Experiment 3 (SD)\*** |  |  |  |  |  |  |  |  |
| Col-0 | 82.2 | 79-86 | 68.6 | 11.9 | 80.5 | 76-86 | 22 | - |
| *ML1:amiRTPPJ V1 #1* | 65.3 | 61-69 | 56.7 | 9.9 | 66.5 | 63-72 | 22 | 1.08E-25 |
| *clv3-10* | 95.8 | 95-101 | NA | NA | NA | NA | 26 | 2.17E-25 |
| *clv3-10;ML1:amiRTPPJ V1 #1_1* | 87.3 | 84-92 | NA | NA | NA | NA | 24 | 4.39E-09 |
| *clv3-10;ML1:amiRTPPJ V1 #1_3* | 85.8 | 82-91 | NA | NA | NA | NA | 23 | 4.05E-05 |
| **Only presented as supplemental material:** |  |  |  |  |  |  |  |  |
| **Experiment 2 (LD)** |  |  |  |  |  |  |  |  |
| Col-0 | 22.8 | 22-25 | 12.1 | 3.2 | 15.3 | 14-16 | 20 | - |
| *ML1:amiRTPPJ V1 #1* | 20.0 | 17-21 | 9.0 | 3.1 | 12.1 | 10-13 | 20 | 4.33E-10 |
| *ML1:amiRTPPJ V2 #5* | 21.0 | 20-22 | 10.3 | 2.9 | 13.2 | 10-14 | 20 | 9.95E-06 |
| **Experiment 3 (SD)\*** |  |  |  |  |  |  |  |  |
| *clv3-7* | 87.5 | 85-91 | NA | NA | NA | NA | 25 | 1.38E-09 |
| *clv3-7;ML1:amiRTPPJ V1 #1_2* | 74.6 | 70-78 | NA | NA | NA | NA | 24 | 5.50E-22 |
| *clv3-7;ML1:amiRTPPJ V1 #1_12* | 76.9 | 73-80 | NA | NA | NA | NA | 24 | 1.47E-19 |
| **Experiment 4 (LD)** |  |  |  |  |  |  |  |  |
| Col-0 | 26.5 | 25-29 | 20.6 | 5.1 | 25.7 | 24-29 | 31 | - |
| *35S:amiRTPPJ V1 #1* | 24.3 | 22-26 | 17.8 | 4.6 | 22.3 | 21-25 | 32 | 1.16E-08 |
| *35S:amiRTPPJ V1 #2* | 23.0 | 21-26 | 16.8 | 4.3 | 21.2 | 19-23 | 32 | 2.24E-11 |
| *tppj* | 25.8 | 24-28 | 19.8 | 4.8 | 24.6 | 22-27 | 35 | 0.04 |
| **Experiment 5 (SD)** |  |  |  |  |  |  |  |  |
| Col-0 | 65.1 | 61-68 | 63.7 | 10.1 | 73.7 | 70-76 | 30 | - |
| *35S:amiRTPPJ V1 #1* | 60.4 | 57-62 | 58.4 | 9.7 | 68.1 | 63-71 | 30 | 6.63E-12 |
| *35S:amiRTPPJ V1 #2* | 58.3 | 56-60 | 57.8 | 9.5 | 67.2 | 66-69 | 30 | 1.47E-17 |
| *tppj* | 59.3 | 57-62 | 58.4 | 9.4 | 67.8 | 65-72 | 30 | 6.15E-15 |
| **Experiment 6 (SD)** |  |  |  |  |  |  |  |  |
| Col-0 | 72.4 | 69-75 | 67.9 | 9 | 77.2 | 74-81 | 26 | - |
| *spl3* | 73.4 | 71-75 | 68.3 | 10 | 78.0 | 75-81 | 23 | 0.15 |
| *spl4* | 74.0 | 71-77 | 69.2 | 14 | 83.0 | 80-86 | 26 | 5.00E-05 |
| *spl5* | 74.0 | 71-77 | 68.0 | 10 | 77.8 | 74-82 | 24 | 0.45 |
| *spl34* | 77.9 | 76-80 | 74.1 | 17 | 90.9 | 87-94 | 24 | 5.04E-09 |
| *spl35* | 73.2 | 71-75 | 68.4 | 10 | 77.9 | 75-81 | 24 | 0.02 |
| *spl45* | 76.8 | 74-78 | 73.0 | 16 | 76.8 | 86-93 | 24 | 1.25E-11 |
| *spl345* | 84.1 | 82-87 | 79.6 | 19 | 99.0 | 94-112 | 27 | 2.79E-26 |

Please note that data marked with an asterisk (\*) were obtained from the same experiment.

**Table S2.**
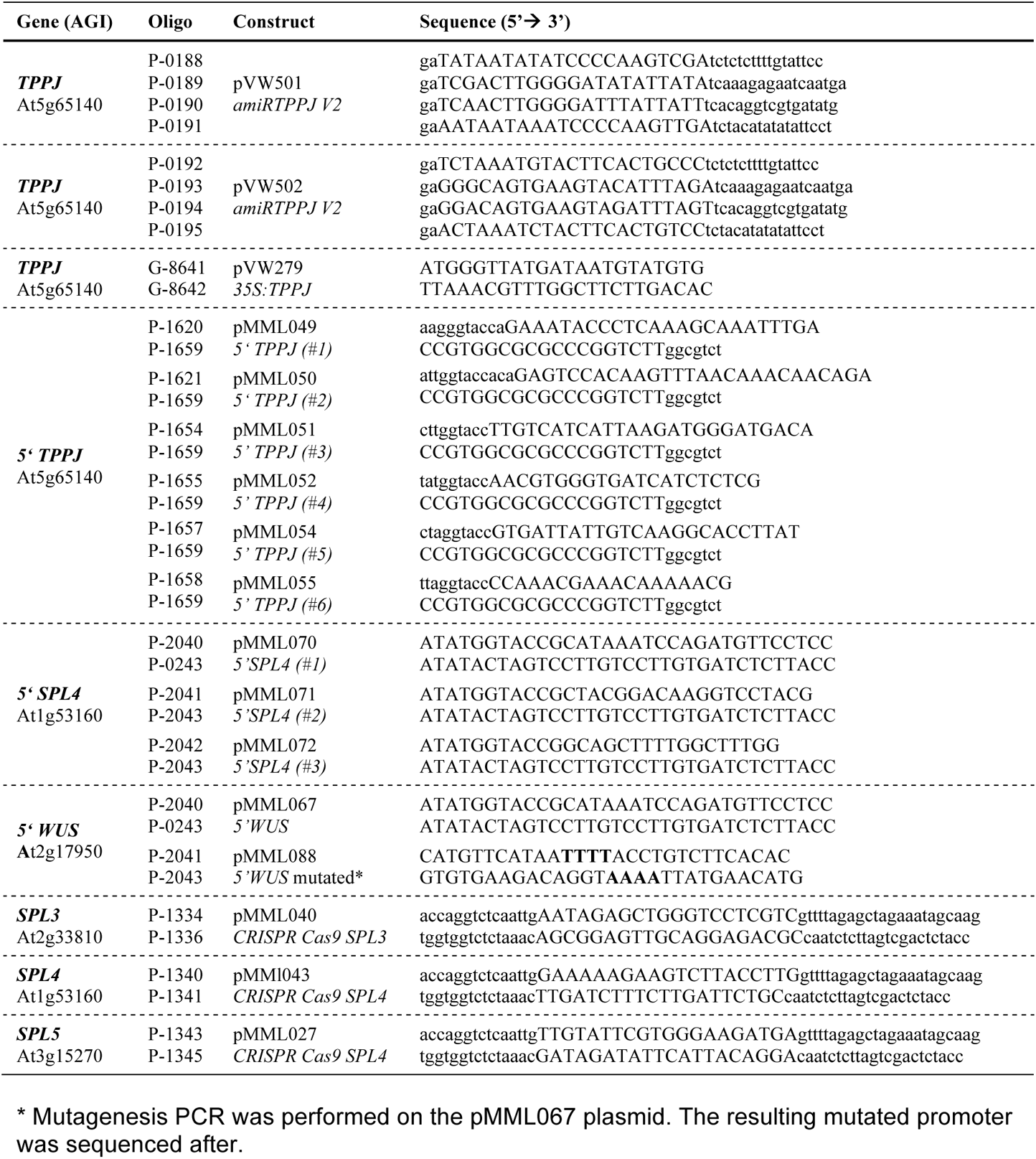
Oligonucleotides for constructs used for the generation of transgenic lines and for transactivation assays.

**Table S3.**
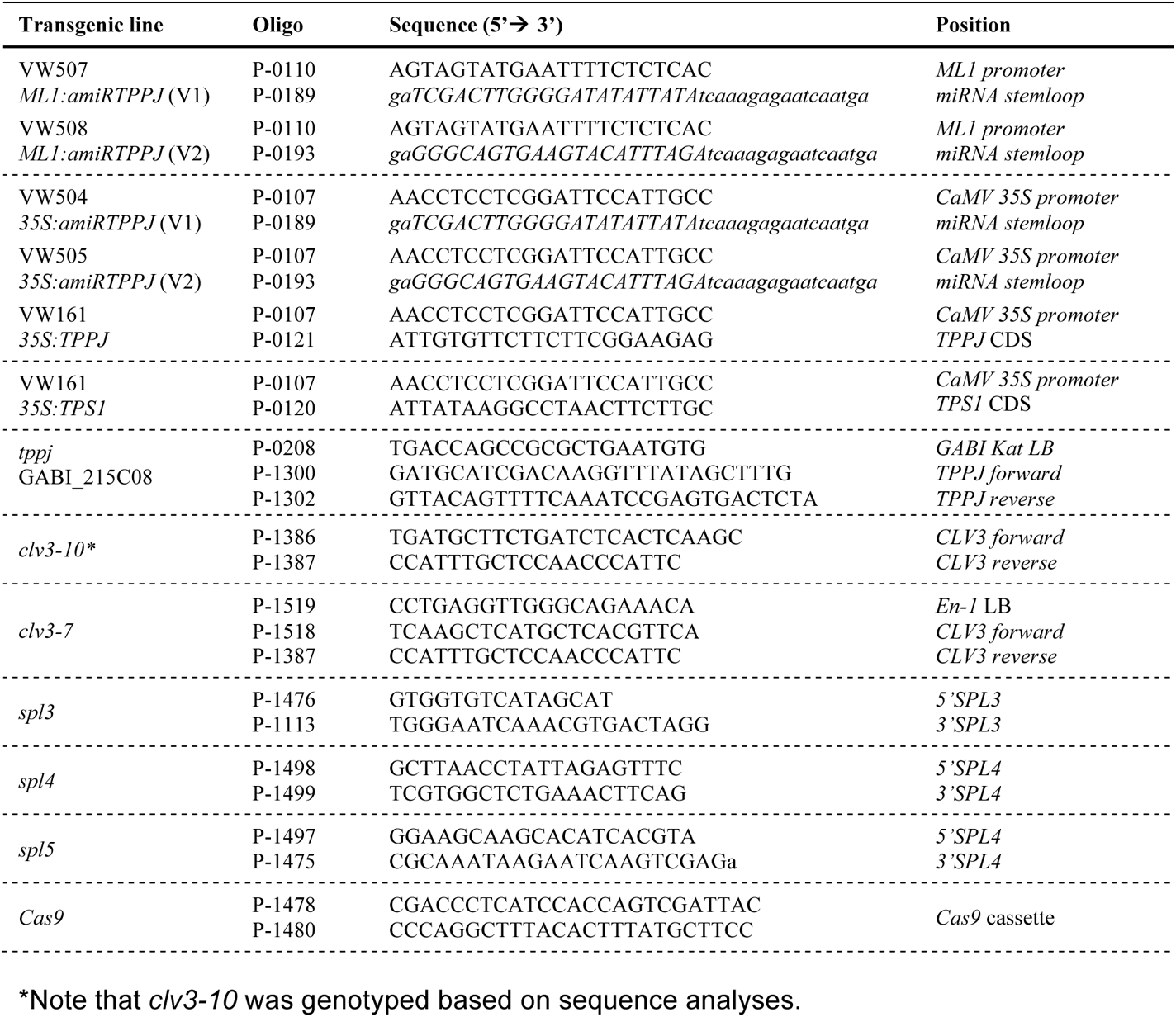
Oligonucleotides used for genotyping.

**Table S4.**
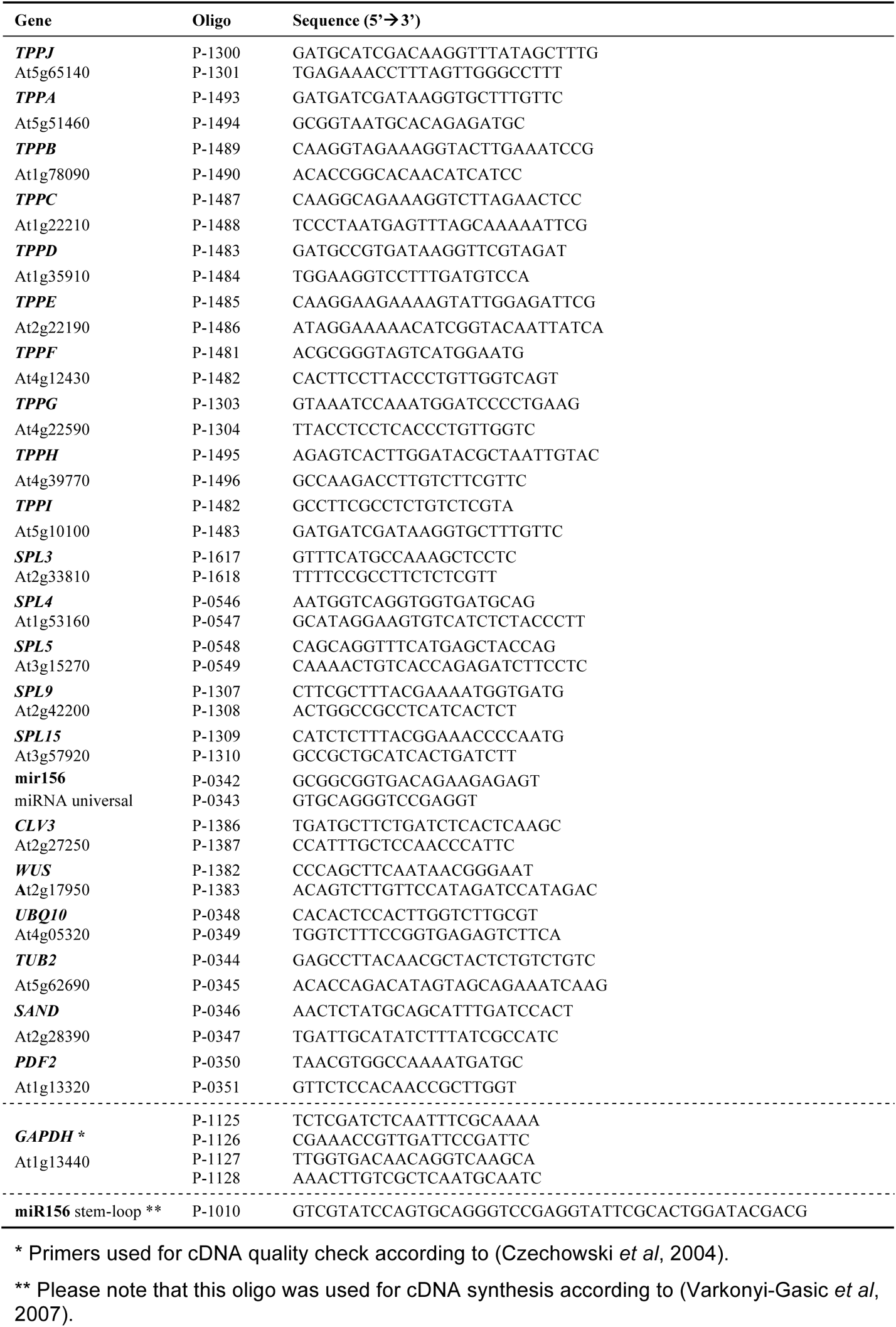
Oligonucleotides used for RT-qPCR.

**Table S5.**
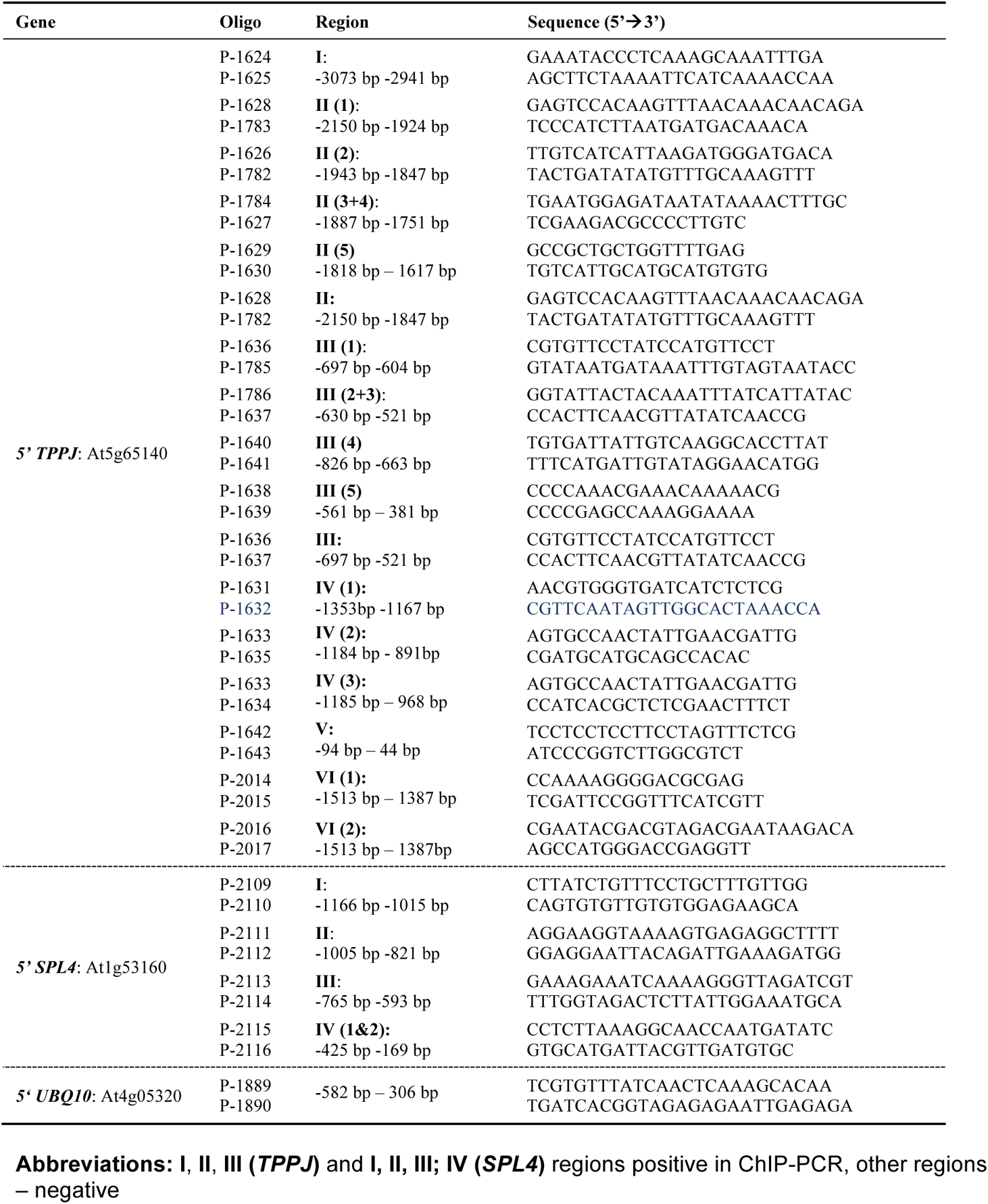
Oligonucleotides used for ChIP-PCR.

**Table S6.**
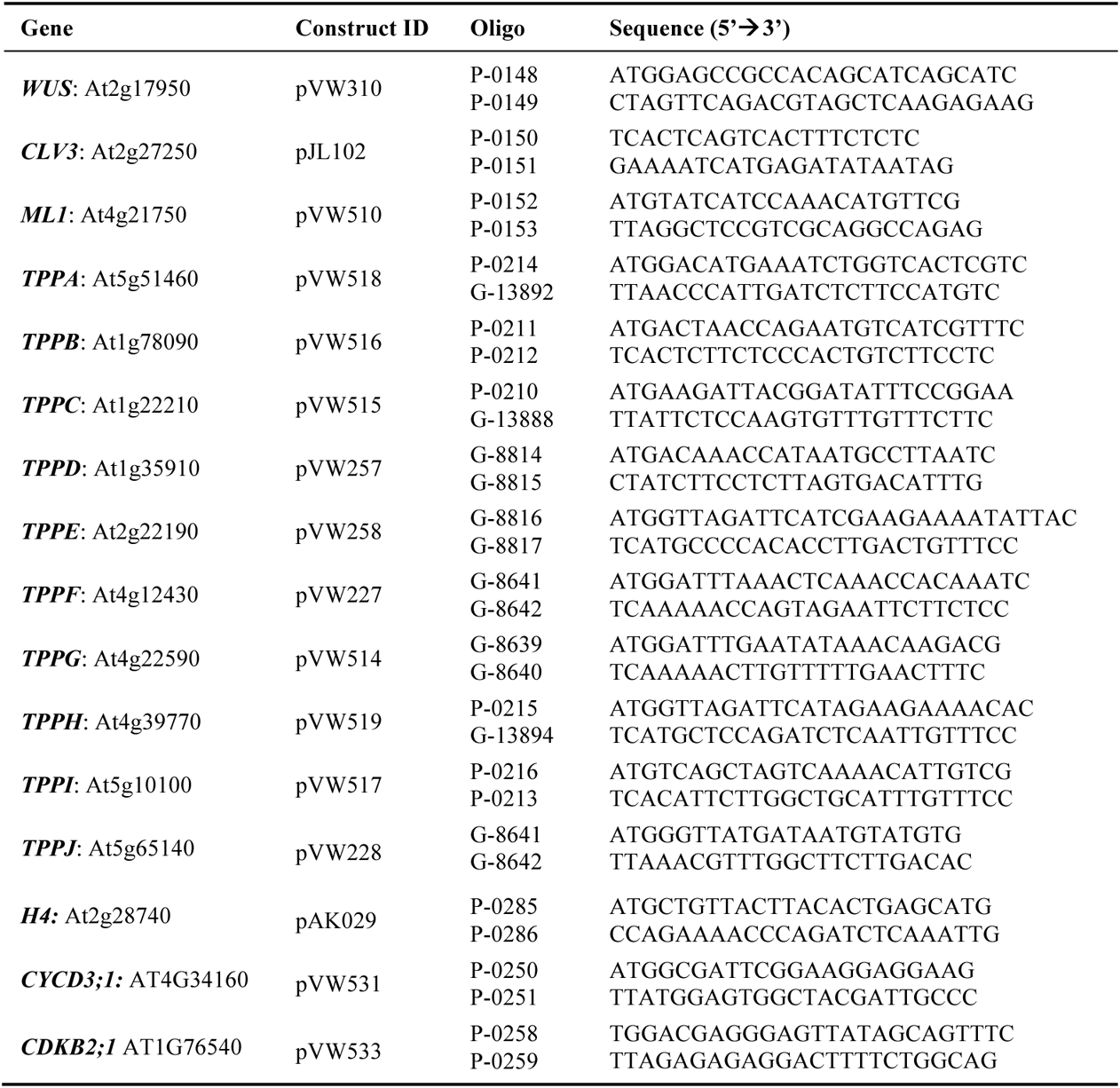
Oligonucleotides used to prepare constructs for probe synthesis.

**Table S7.**
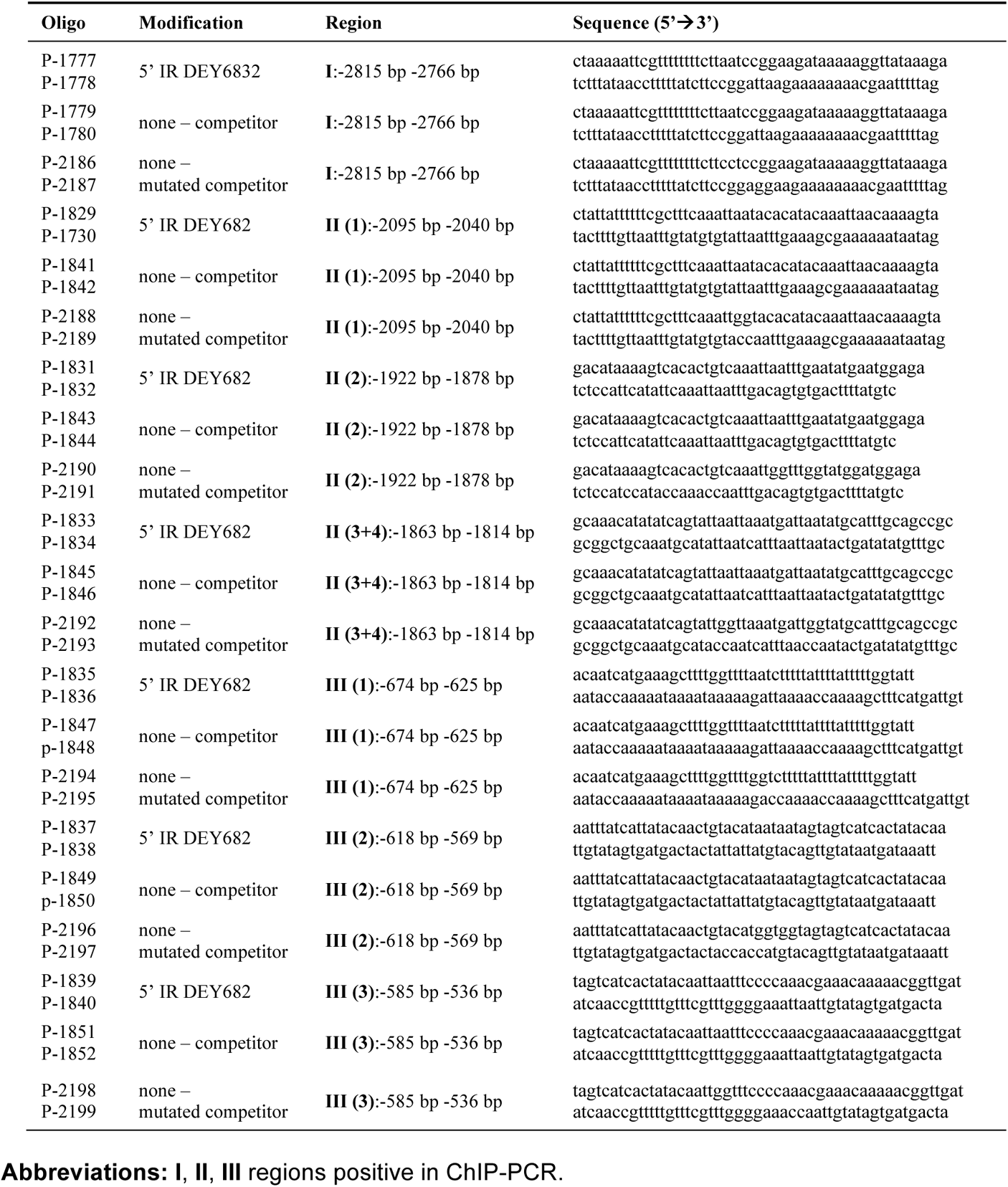
EMSA oligonucleotides (probes) for *5’TPPJ* At5g65140.

**Table S8.**
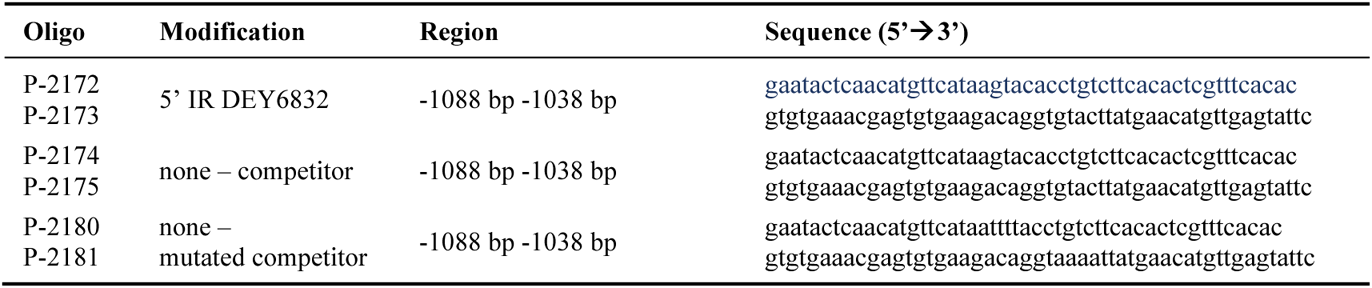
EMSA oligonucleotides (probes) for *5’WUS* At2g17950.

